# Ontogeny of the vagal gut-brain axis

**DOI:** 10.1101/2025.05.01.651736

**Authors:** Kendall J. Lough, Jackson C. Knox, Sheryl S. Moy, Juliet S. King, Scott E. Williams

## Abstract

Gut-brain communication is a key component of homeostasis which regulates behaviors such as appetite and reward. Intestinal entero-endocrine cells (EECs) translate nutrient intake into signals which affect sensation and behavior, in part through synapse-like contacts with vagal nodose neurons. This direct neuroepithelial circuit regulates feeding *(1–3)*, yet its genesis and role in higher order behaviors remains unknown. We find that EECs first contact nodose neurons in utero, and these interactions require brain-derived neurotrophic factor (BDNF). We show that BDNF regulates the dynamics of these interactions, which underlie EEC-evoked optogenetic excitation of the circuit. In addition to feeding, we discover that this circuit regulates sociability and inhibitory behavioral control. These studies define the ontogeny of a direct gut-brain circuit responsible for early physiology and behavior.

---

Neuroepithelial sensory circuits are a key feature of environment-facing epithelia, which feed mechanical and chemical information back to the brain, allowing organisms to interpret and interact with their surroundings. This is true of the epidermis (*4*), lungs (*5*), and the intestine, the latter of which relies on epithelial sensory cells called enteroendocrine cells (EECs) to translate luminal information into cellular signals that modify neuronal activity (*6*). The gut lumen hosts a complex chemical environment, where dietary nutrients, microbial products, drugs, and noxious agents can act upon the absorptive epithelia to alter digestion, behavior, and nociception. EECs express surface receptors for many of these compounds, and in turn release hormones and neuropeptides to mediate changes in sensation and physiology. Classically, EECs are thought to signal to the CNS via endocrine signaling, but recent studies have provided evidence that EECs can also directly interact with afferent projections and drive neuronal activity on synapse-like timescales (*1, 7*).

The intestine receives sensory afferent innervation from the nodose and dorsal root ganglia (DRG), via the vagal and sacral nerves, respectively (*8*). While both afferents can regulate satiety (*1, 9*), DRGs may uniquely mediate nociception in the gut (*10*). These variant roles in physiology may correlate with the EEC subpopulations these fibers innervate: While glutamatergic, cholecystokinin (CCK)-expressing EECs rely on vagal nodose afferents to influence behavior, serotonergic enterochromaffin cells require intact sacral fibers emanating from DRGs (*9*).

These biases suggest that EECs leverage unique molecular pathways to matchmake with specific neurons. However, the ontogeny of these circuits has never been described, and whether direct connections regulate specific behaviors remains unknown. Here, we investigated the formation and role of the EEC-nodose circuit using a variety of transgenic mouse models, rabies trans-synaptic tracing, single-cell transcriptomic, and optogenetic tools, including novel techniques for *in utero* intestinal delivery of viruses and *ex vivo* live imaging of gut-ganglion co-cultures. We identify BDNF as essential for circuit formation and establish the EEC-nodose circuit as a mediator of not only satiety, but also sociability and anxiety. Collectively, these studies define the origins of direct gut-brain communication, and support human studies that link *BDNF* and its receptor *NTRK2* to childhood obesity and autism-spectrum disorder (*11–13*).

## Intestinal neural-epithelial interactions form during embryogenesis

EECs form connections with synapse-like properties to vagal afferents from the nodose ganglion, often through a basal, axon-like process called a “neuropod” (*1*). To understand how and when these connections form, we utilized a genetic, cell type-specific membrane-labeling strategy to mark embryonic EECs and analyze their proximity to neuronal processes (fig. S1A). We crossed *Neurod1*^*Cre*^—expressed by all terminally differentiated EECs (*14*)— with the *Rosa*^*mTmG*^ reporter (*15*), resulting in membrane anchored eGFP (mGFP) expression specifically in all committed EECs (Fig. 1A,B).

**Figure 1.**
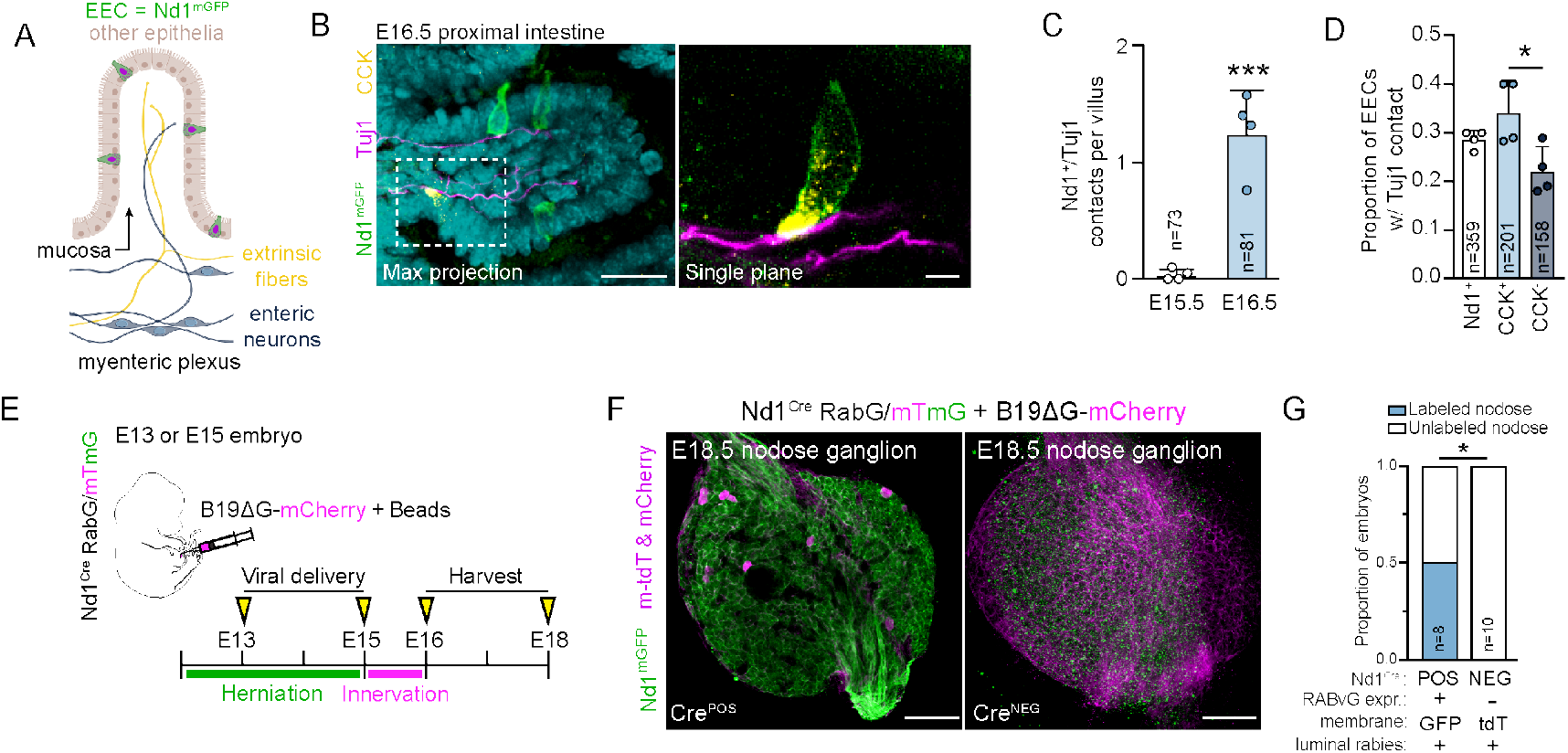
Embryonic enteroendocrine cells interact with nodose neurons at E16.5. (A) Depiction of EEC genetic labeling strategy. *Neurod1*^*Cre*^; *Rosa*^*mTmG*^ is used to label EECs with membrane (m)-GFP (*Nd1*^*mGFP*^), which exclusively reside within the epithelia (beige) lining intestinal villi. Beneath the epithelia, neural fibers project into the mucosa. These can originate from either extrinsic neurons (yellow; e.g. vagal nodose neurons) or enteric neurons (blue), whose soma reside within the submucosal or myenteric plexuses in the gut wall. (B) Max projection (left) and single plane image of boxed region (right) in E16.5 *Nd1*^*mGFP*^ proximal intestine stained for mGFP (green), Tuj1 (magenta), and CCK (yellow). (C) Quantification of frequency of Nd1+ cells contacting Tuj1+ neural fibers per villus from E15.5 and E16.5 proximal intestines. Data are from 4 individuals (dots) and at least 2 separate litters; *n* indicates total villi assessed. (D) Proportion of Nd1+ EECs with Tuj1+ contacts. Each dot is data from one individual embryo; *n* refers to the number of individual EECs. (E-G) Rabies transsynaptic tracing experiments. (E) Experimental design where B19ΔG-mCherry rabies is injected into the umbilical hernia of E13 or E15 embryos (prior to mucosal innervation) where it is able to infect *NeuroD1*^*Cre*^+ EECs which express the RabG protein necessary for trans-synaptic tracing. (F) Max projection images of whole nodose ganglia from E18.5 *Nd1*^*RABvG/mGFP*^ embryos (left) or Cre-negative *Nd1*^*mRFP*^ controls (right) with luminal intestinal delivery of B19ΔG-mCherry rabies virus. Rabies-derived mCherry+ nodose labeling indicates likely transfer from intestinal viral payload and is only observed in *Neurod1*^*Cre+*^ embryos. Note that Nd1+ nodose fibers appear mT+ (right) rather than mG+ (left), confirming the absence of Cre in these controls. (G) Quantification of the proportion of embryos with mCherry nodose labeling in *Neurod1*^*Cre*^ POS and NEG embryos. Scale Bars, 25 µm (A, left panel), 5 µm (A, right), 100 µm (F). * *P* < 0.05, *** *P* < 0.001, by student’s t-test.

Among *Neurod1*^*Cre*^; *Rosa*^*mTmG*^ mGFP+ cells (hereafter referred to as *Nd1*^*mGFP+*^), both substance P-positive (SubP; *Tac1*) enterochromaffin cells (EC) and cholecystokinin-enriched (CCK; *Cck*) EECs (*9, 16*) could be detected as early as embryonic day (E)15.5. Expression of both maturation markers increased from E15.5-E17.5 and were most abundant in the proximal intestine at all ages (fig. S1B-G). To determine if the tempo of mucosal innervation parallels EEC specification, we labeled axons with the pan-neuronal marker Tuj1 (*Tubb3*). We detected neuronal projections in the mucosa of nearly all villi of the proximal small intestine at E16.5 but not at E15.5, when neurites were restricted to the underlying myenteric plexus (fig. S1H-J) (*17*). We observed frequent instances of overlap between mGFP+ EEC-membranes and Tuj1+ axons in E16.5 proximal intestines (Fig. 1B), indicating that some embryonic EECs may make contact with neuronal projections. While instances of EEC-neuron contact were observed in most villi at E16.5 (50/81 villi; 61.7%) and single villi often had multiple EECs with Tuj1+ contact (Fig. 1C), such contacts were extremely rare at E15.5 (2/73 villi; 2.7%), suggesting that this timepoint is a key developmental milestone in the formation of gut-brain communication. Furthermore, these contacts were enriched among CCK+ compared to CCK-EECs (Fig. 1D), suggesting that neurons may preferentially interact with specific EEC subtypes.

## The enteroendocrine cell-nodose circuit forms *in utero*

We next sought to determine if the mucosal projections observed at E16.5 might arise from extrinsic nodose or DRG neurons. To do so, we relied on two Cre alleles: *Neurod1*, expressed by nodose neurons (fig. S2A), and Vesicular glutamate transporter-2 (*Vglut2*), expressed by both nodose (fig. S2B) and DRG neurons (*8*). DRGs can further be distinguished by expression of the neurotrophic receptor TrkA (*Ntrk1*) (*8*), which is largely absent in the nodose (fig. S2C). While both *Nd1*^*mGFP*+^ and *Vglut2*^*mGFP*+^ projections could be observed in E16.5 proximal mucosa (fig. S2D,E) (*8*), TrkA+ fibers were only detected in the myenteric plexus (fig. S2F), as *Vglut2*^*mGFP*+^ projections in the mucosa were always TrkA-negative (fig. S2G). Finally, Phox2B+ nuclei of enteric neurons located in the gut wall were never *Nd1*^*mGFP*+^ (fig. S2H). Collectively, these data suggest that afferents emanating from the nodose ganglia are likely the only source of extrinsic sensory innervation in the proximal intestinal mucosa at E16.5.

To determine whether embryonic EECs contact nodose neurons, we devised an *in utero* strategy to perform transsynaptic tracing from the intestine using modified rabies virus. The SADB19-G strain of rabies virus (B19ΔG) displays a natural tropism for EECs (*1*), but is incapable of spreading across neuronal junctions because it lacks the G glycoprotein. Therefore, we crossed *Neurod1*^*Cre*^ with an inducible G glycoprotein allele (*RABvG*; *Nd1*^*RABvG*^), thereby allowing rabies-infected EECs to spread the virus to associated neurons (*18*). We leveraged the window of embryonic umbilical herniation to deliver viral payloads to the midgut—in this case, B19ΔG rabies virus expressing an mCherry reporter mixed with fluorescent microbeads—via ultrasound-guided microinjection (Fig 1E). In half of injected embryos, fluorescent beads could be observed throughout the gut lumen, confirming successful intraluminal delivery of the rabies virus (fig. S3A,B). Cytoplasmic accumulation of the mCherry reporter was only observed in nodose neurons of *Neurod1*^*Cre*^-positive embryos, and never seen in *Neurod1*^*Cre*^-negative controls (Fig. 1F,G). These data confirm that rabies virus can transfer from *Neurod1*^*Cre*^-positive EECs to neurons in the nodose ganglion, suggesting they synapse with vagal nodose afferents.

## Single-cell profiling of embryonic EECs reveals *Bdnf* as a candidate factor

We next sought to identify candidate factors which might influence the formation of EEC-nodose connections using single-cell RNA sequencing (scRNAseq) of FACS-isolated GFP+ EECs from E15-E17 *Nd1*^*mGFP*^ embryonic small intestines (fig. S4A). Unbiased clustering analysis identified 17 distinct populations with divergent transcriptomic profiles (fig. S4B). We defined population identity by marker gene expression (fig. S4C), segregating epithelia via expression of *Epcam, Cdh1*, and *Vil1* (fig. S4D). This included EECs, one unidentified secretory population (Sec1) and three non-secretory epithelia, including *Fabp1*+ enterocytes (Fig. 2A). EEC subtypes were defined by gastric hormone expression, with the enterocyte (EC) population used as a negative control (Fig. 2B). Embryonic EECs were enriched for classic gastric hormones, as well as receptors for macronutrients (fig. S4F), and many pre- and post-synaptic markers (fig. S4G), suggesting that embryonic EECs already harbor the necessary equipment to perform their chemosensory signaling function.

**Figure 2.**
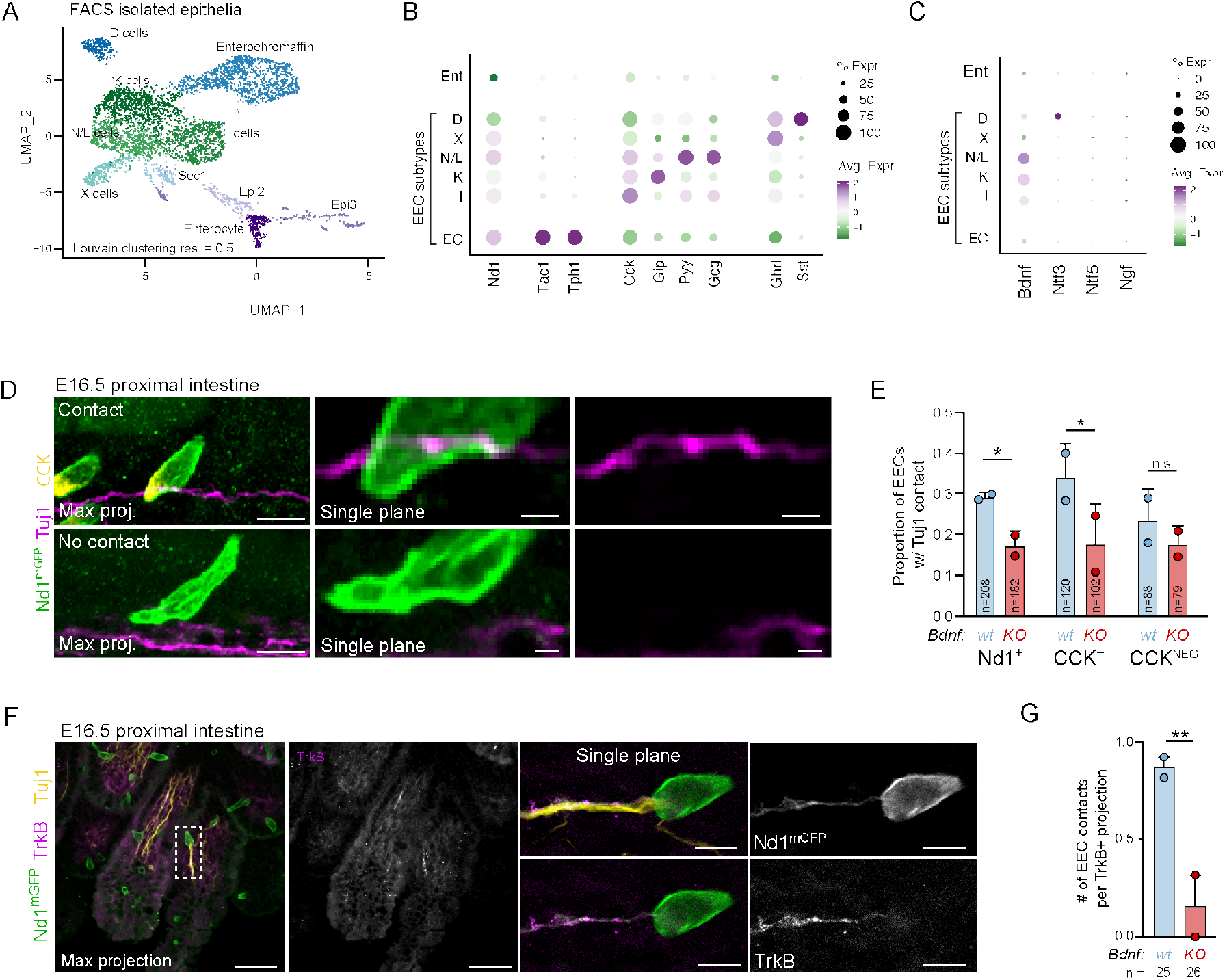
EEC-derived *Bdnf* regulates neuronal interactions. (A) Uniform manifold approximation and projection (UMAP) of FACS-isolated and sequenced epithelia including 3,791 embryonic (E15-17) EECs sorted into 7 distinct clusters, in addition to 3 clusters of non-secretory epithelia. (B) Dot plot of expression of gastric hormones across EEC clusters, with enterocytes as negative control. Here and in all dot plots, green hues represent low expression and purple hues high expression, while the size of the dot indicates the proportion of positive cells in the cluster. (C) Dot plot of neurotrophin expression across EEC clusters. *Bdnf* is uniquely expressed by a subset of I, K, and N/L cells. Ent = enterocytes, EC = enterochromaffin cells. (D) Image series—max projection (left), single plane (middle) and single channel (right)—of *Nd1*^*mGFP*^ EECs from E16.5 proximal intestine stained for mGFP (green), Tuj1 (magenta) and CCK (yellow), depicting instances of EEC-neuron contact (top series) or no contact (bottom series). (E) Quantification of proportion of EECs interacting with Tuj1+ axons in E16.5 proximal intestine of *Bdnf*^*+/+*^ (WT, blue) and *Bdnf*^*Nd1-cKO*^ (KO, red) embryos. Dots represent data from individual embryos, from two separate litters; *n* refers to number of EECs. (F) Wholemount images from E16.5 proximal intestine of *Nd1*^*mGFP*^ embryo stained for mGFP (green), TrkB (magenta) and Tuj1 (yellow). Left panel is a max projection of multiple villi. Middle and right panels are magnification of single EEC and axon from left panel displaying contact between EEC and TrkB+ axon. (G) Quantification of the number of EEC interactions per TrkB+ axon from E16 and E17 proximal intestine. Dots represent data from individual embryos (one WT and one KO embryo from two separate litters); *n* refers to number of axons. Scale bars, 10 µm (D left panels, F right panels), 2 µm (D right panels), 50 µm (F left panels). *P* values determined by student’s unpaired t-test; ** *P* < 0.01. * *P* < 0.05. *n*.*s*. = not significant.

We combed EEC transcriptomes for genes related to circuit formation or function, leading us to identify Brain-derived neurotrophic factor (*Bdnf*) (Fig. 2C, fig. S4E). BDNF is one of four neurotrophins, secreted ligands that regulate the development of neural circuits (*19, 20*). BDNF binds exclusively to tyrosine receptor kinase B (TrkB or *Ntrk2*) (*21*), which promotes synaptogenesis, encouraging sustained interaction between the ligand-source and receptor-expressing neuron (*22*). Our data indicate that CCK-expressing EECs are uniquely enriched for *Bdnf* (Fig. 2C). Fittingly, nodose neurons are enriched for TrkB, and BDNF is essential for growth and survival of nodose neurons (*23*). In contrast, DRGs predominantly express the mismatched receptor (TrkA), while enteric neurons express TrkA and TrkC, but not TrkB (*24*). These data support a model wherein EEC-derived BDNF biases EECs to form interactions with TrkB+ nodose neurons.

## EEC-derived BDNF regulates neural-epithelial interactions, *in vivo*

To test this hypothesis, we first confirmed that a subset of embryonic EECs express *Bdnf* at E16.5 (fig. S5A) by crossing a *Bdnf*^*2A-Cre*^ mouse with the *Rosa*^*mTmG*^ reporter (*Bdnf*^*2A-Cre*^,*Rosa*^*mTmG*^; i.e. *Bdnf*^*mGFP*^). *Bdnf*^*mGFP+*^ EECs formed close associations with mucosal projections (fig. S5B), and E16.5 nodose neurons are enriched for TrkB (fig. S5C). Next, we crossed *Nd1*^*mGFP*^ animals with a *Bdnf*^*fl*^ strain (*25*) to selectively knock out *Bdnf* in *Neurod1*-expressing EECs (*Bdnf*^*Nd1-cKO*^). We evaluated E16.5 mGFP+ EECs for the presence or absence of neuronal contact (Fig. 2D) and found that loss of *Bdnf* selectively reduced the frequency of this interaction in CCK+ EECs (Fig. 2E). The lack of a strong effect on CCK^Neg^ EECs fits with our scRNAseq data, which show that *CCK, Gip*, and *Pyy* are the primary hormones co-expressed with *Bdnf* (Fig. 2B,C). To define how *Bdnf* loss impacts nodose projections, we assessed how frequently TrkB+/Nd1^mGFP^ axons interacted with EECs (Fig. 2F). In wild-type animals, these projections frequently interacted with one or more EECs, while EEC-associations were extremely rare following *Neurod1*^*Cre*^ driven *Bdnf* deletion (Fig. 2G). These findings suggest that BDNF regulates EEC-neuron interactions and is particularly essential for associations between CCK+ EECs and TrkB+ nodose neurons.

## An *ex vivo* co-culture model to study extrinsic innervation of the small intestine

To better understand how *Bdnf* regulates EEC-nodose interactions, we engineered an explant co-culture system to enable dynamic visualization of intestinal innervation by nodose neurons (fig. S6A). Contact-naïve E15.5 intestines and/or nodose ganglia were cultured *ex vivo* and imaged daily for 7 days. *Nd1*^*mGFP*^ intestines were cut longitudinally to expose the lumen yet remained viable and maintained their villus like architecture (fig. S6B). Importantly, no *Nd1*^*mGFP*^ neuronal fibers were observed in isolated *Nd1*^*mGFP*^ intestinal explants or the intestines of freshly placed co-cultures (fig. S6C), confirming that mGFP+ projections observed in *Nd1*^*mGFP*^ intestines, *in vivo*, must come from an extrinsic source. When a *Vglut2*^*mGFP*^ nodose ganglion was co-cultured adjacent to *Nd1*^*mGFP*^ intestine (Fig. 3A), mGFP+ neuronal fibers could be observed growing into the intestinal sample after several days (Fig. 3B; fig. S6D), eventually contacting *Nd1*^*mGFP*^ EECs (fig. S6E). These data confirm that embryonic EECs can form direct interactions with nodose neurons and establish a new model system to evaluate the process of intestinal circuit formation.

**Figure 3.**
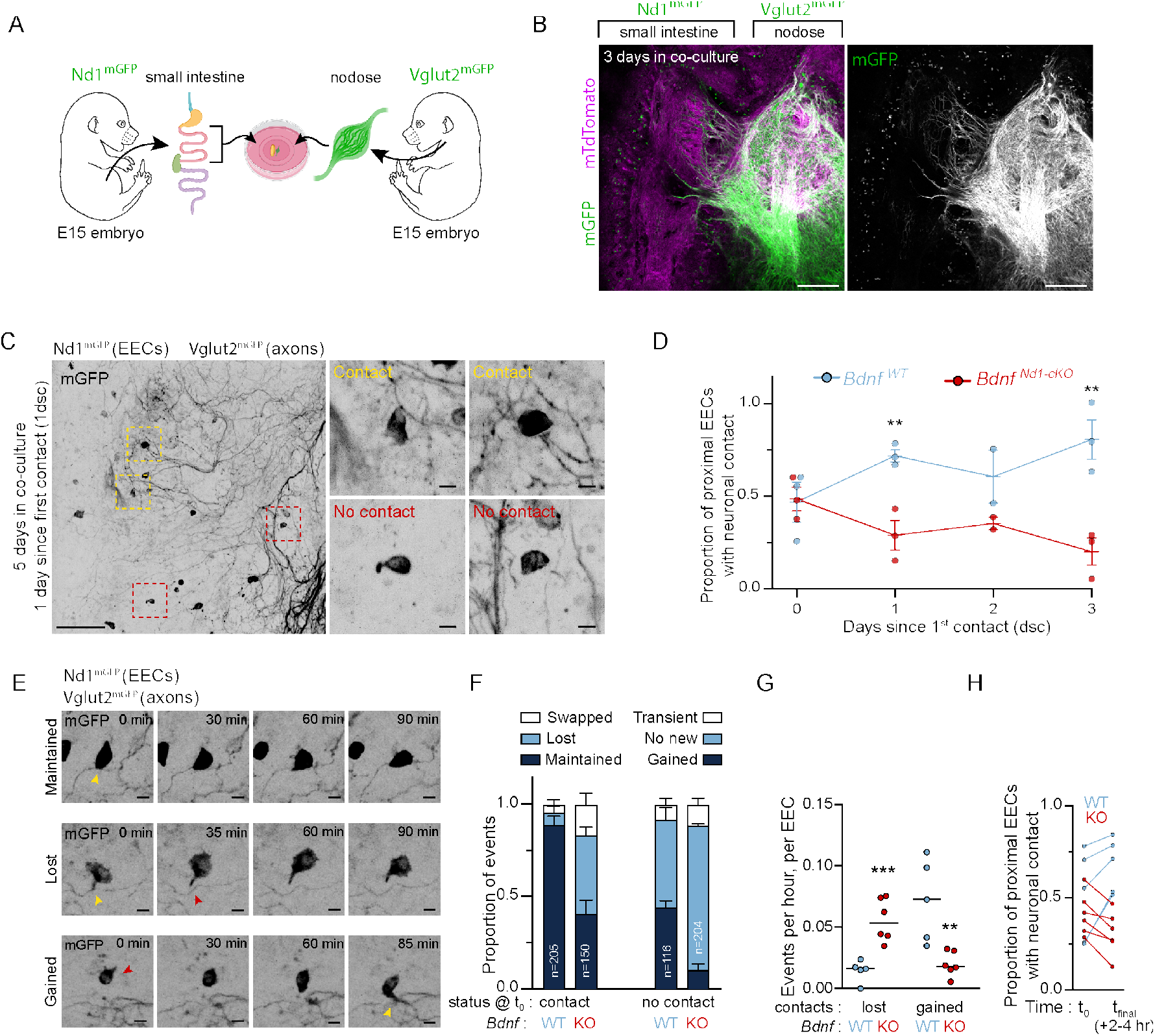
*Ex vivo* co-culture model recapitulates EEC-nodose interactions. (A,B) Schematic outline (A) and image still (B) of 3 day mGFP labeled gut-nodose explant co-culture live imaging experiment. In this paradigm, EECs in the intestinal explant are *Nd1*^*mGFP+*^ while projections from the nodose are *Vglut2*^*mGFP+*^. Cre-negative cells are mTdTomato+. Single channel (mGFP) shown at right. (C) Movie stills from co-cultures after 5 days in co-culture, one day after first observed EEC-neuron contact (1dsc). Low magnification left panel depicts numerous *Nd1*^*mGFP*^ EECs in close proximity to *Vglut2*^*mGFP*^ nodose projections. Boxed regions show higher magnification examples of contacts (top) or no contact (bottom) shown at right. (D) Quantification of proportion of proximal EECs (<25 µm from a nodose axon) with nodose interactions. *Bdnf*^*WT*^ (WT, blue) EECs increasingly interact with nodose neurons over time, while *Bdnf*^*Nd1-cKO*^ (KO, red) EECs decrease the proportion with contacts. (E) Movie stills from 1dsc co-cultures, depicting examples of dynamic interaction behaviors: contact maintained (top), contact lost (middle) or contact gained (bottom); yellow arrowheads indicate site of contact, while red arrowheads indicate absence of contact. (F) Quantification of dynamic EEC-nodose interactions (0-2dsc) as a proportion of total events, comparing *Bdnf*^*WT*^ and *Bdnf*^*Nd1-cKO*^ EEC behavior, categorized by whether there was initial contact (left) or not (right). *Bdnf*^*Nd1-cKO*^ EECs were more likely to swap or lose contact and less likely to gain contacts. Data depicted are from 3 biological and technical replicates; *n* refers to the total number of EECs. (G,H) Quantification of the number of contacts lost or gained for *Bdnf*^*WT*^ (blue) and *Bdnf*^*Nd1-cKO*^ (red) EECs. Data are shown as events per hour across all imaging sessions for 0-2dsc (G), and proportion of EECs with contact at the beginning and end of a 2-4h experiment (H). Each dot represents an imaging experiment. Scale bars, 300 µm (B), 100 µm (C left panel), 10 µm (C right panels, E). *P* values determined by student’s unpaired t-test. ** *P* < 0.01, *** *P* < 0.001.

## BDNF regulates dynamic enteroendocrine-nodose interactions

To specifically query the function of BDNF in EECs, we paired WT or *Neurod1*^*Cre*^; *Bdnf*^*fl/fl*^ (*Bdnf*^*Nd1-cKO*^) intestines with wild-type nodose explants. *Nd1*^*mGFP*^ intestines were isolated from *Bdnf*^*+/+*^ (*Bdnf*^*WT*^) or *Bdnf*^*Nd1-cKO*^ embryos and co-cultured alongside *Vglut2*^*mGFP*^ nodose ganglia, where we evaluated the proportion of EECs contacting neuronal fibers. Loss of *Bdnf* did not alter the proportion of EECs contacting neurons on the first day contact was observed (0 days since contact, or 0 dsc), suggesting that BDNF does not participate in long-range axon guidance. However, while *Bdnf*^*WT*^ EECs increasingly interacted with nodose fibers over subsequent days, *Bdnf*^*Nd1-cKO*^ EECs decreased their rate of contact (Fig. 3C,D), confirming that BDNF is needed to form and stabilize EEC-nodose interactions.

When observed across shorter time periods, we documented a variety of EEC-neuron behaviors (Fig. 3E, Movie S1-4). EECs with established contacts either maintained, lost, or swapped contacts over time. EECs without initial contacts formed new, sustained contacts, transient interactions, or failed to form any contact (“no new”). *Bdnf*^*WT*^ EECs predominantly maintained initial contacts, while *Bdnf*^*Nd1-cKO*^ EECs frequently lost or swapped their interactions (Fig. 3F). Furthermore, while ∼40% of *Bdnf*^*WT*^ EECs formed new interactions, *Bdnf*^*Nd1-cKO*^ EECs did so <10% of the time (Fig. 3H). Additionally, *Bdnf*^*Nd1-cKO*^ EECs were more likely to lose contacts and less likely to gain new contacts than their wild-type counterparts (Fig. 3G). *Bdnf*^*Nd1-cKO*^ guts consistently reduced the proportion of EECs contacting neurons, while *Bdnf*^*WT*^ samples consistently increased this proportion (Fig. 3H). These data demonstrate that BDNF regulates the frequency and stability of interactions between EECs and proximal nodose projections.

## Direct communication underlies EEC-evoked nodose activity

EECs regulate neuronal activity through a combination of endocrine, paracrine, and direct, contact-dependent mechanisms, but the specific contribution of each modality remains poorly understood. Since *Bdnf* loss specifically mitigated the ability of EECs to sustain contact with nodose neurons, we sought to determine how loss of direct communication affected their ability to excite nodose neurons. Therefore, we co-cultured intestines from *Bdnf*^*WT*^ embryos where expression of channel rhodopsin is restricted to EECs (*Neurod1*^*Cre*^; *Bdnf*^*+/+*^; *LSL-ChR2-YFP* (*26*), i.e., *Nd1*^*ChR2*^ *Bdnf*^*WT*^) alongside nodose ganglia expressing the GCaMP6s fluorescent calcium reporter (*Vglut2*^*Cre*^; *LSL-GCaMP6s* (*27*), i.e., *Vglut2*^*GCaMP6s*^) (fig. S7A), and stimulated EECs to secrete their neuromodulatory cargo (fig. S7B,C). After 4 days in co-culture, blue light stimulation of *Nd1*^*ChR2*^ *Bdnf*^*WT*^ intestines was sufficient to drive fast (sub-second) calcium spikes in many nodose neurons (Fig. 4A). In contrast, stimulation excited very few nodose neurons grown alongside a *Nd1*^*ChR2*^ *Bdnf*^*Nd1-cKO*^ intestinal sample (Fig. 4B). A 1 second stimulation evoked rapid spikes of GCaMP6s fluorescence (>0.25 ΔF/ΔF_MAX_, within 1 second) in 69% of *Nd1*^*ChR2*^ *Bdnf*^*WT*^ co-cultures, but only 29% of nodose neurons cultured alongside *Nd1*^*ChR2*^ *Bdnf*^*Nd1-cKO*^ intestines (Fig. 4C-E). These proportions closely mirror the contact rates observed for *Bdnf*^*WT*^ (72±6%) and *Bdnf*^*Nd1-cKO*^ (29±6%) EECs at 1dsc (Fig. 3D; equivalent to 4-5dcc), suggesting that rapid EEC-evoked activity requires sustained, direct contact with nodose projections. *Vglut2*^*GCaMP6s*^ nodose neurons cultured without an intestine (i.e. “no gut” controls) never responded to stimulation (fig. S7D-F).

**Figure 4.**
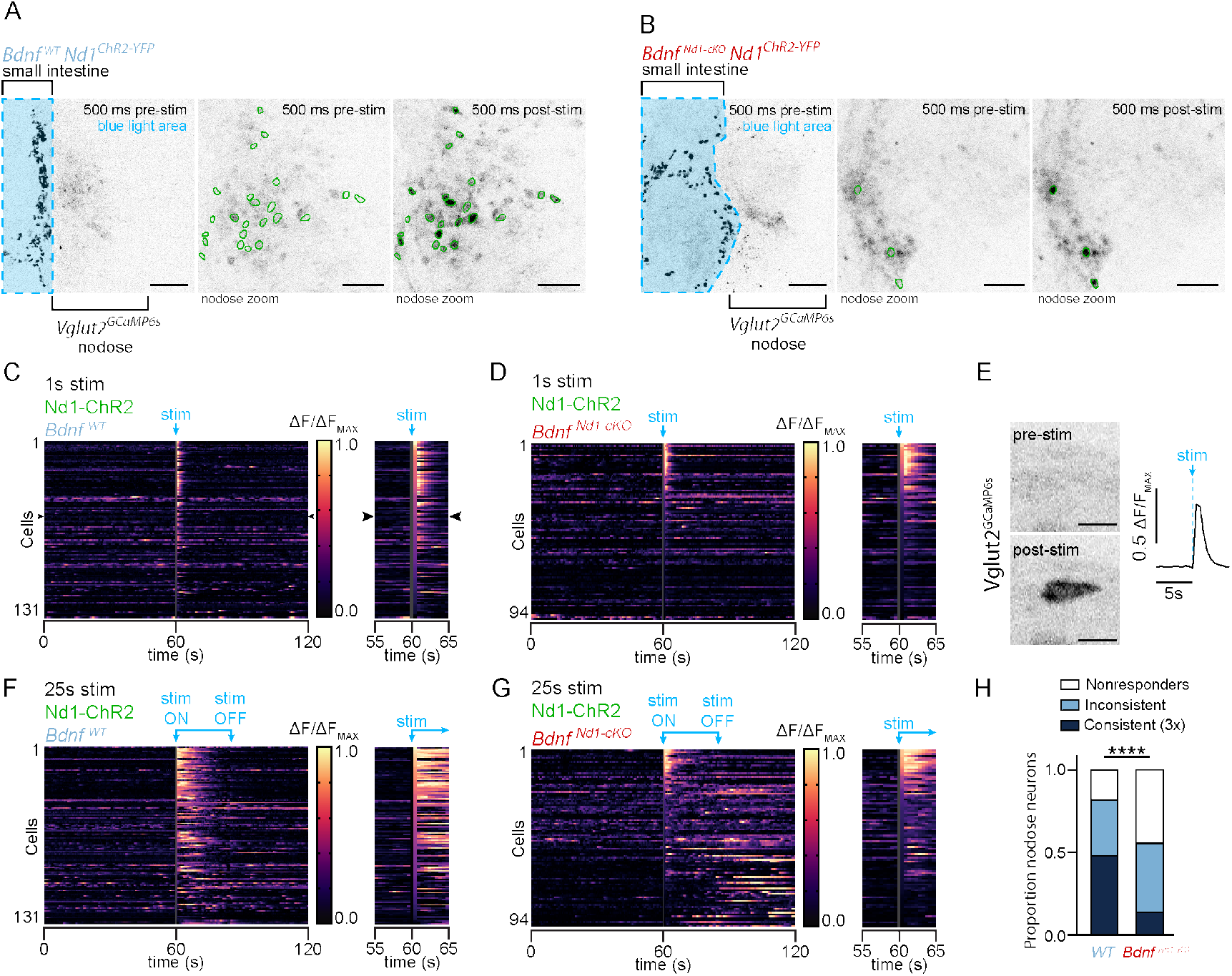
BDNF is required for consistent EEC-evoked excitation of nodose neurons, *ex vivo*. (A,B) Stills from live imaging of 4 day old optogenetic co-culture of *Nd1*^*ChR2*^ intestine adjacent to a *Vglut2*^*GCaMP6s*^ nodose ganglion. (A) *Bdnf*^*WT*^ intestine and (B) *Bdnf*^*Nd1-cKO*^ intestine. Green fluorescence (YFP in intestinal EECs and GCaMP6s in nodose neurons) is shown in inverted greyscale. Left panel shows full co-culture and blue-light stimulation region. Middle and right panels are zooms of nodose ganglion showing GCaMP6s fluorescence immediately prior (middle) or immediately after (right) 1s blue light stimulation. Neurons displaying evoked excitation are outlined in green. (C,D) Heatmap kymograph traces of GCaMP6s fluorescence of individual neurons in *Nd1*^*ChR2*^; *Vglut2*^*GCaMP6s*^ 4 day co-culture experiments with *Bdnf*^*WT*^ intestine (C) or *Bdnf*^*Nd1-cKO*^ intestine (D) with 1s optogenetic stimulation. Each row represents Ca^2+^ fluctuations for an individual neuron, with 2 minutes of activity shown at left, and 10s of activity—immediately before and after stimulation—shown at right. Black arrowheads indicate cell shown in (E). (E) Movie stills and F-F_0_ trace (GCaMP6s fluorescence minus baseline) showing individual neuron (black arrowheads in (C) immediately prior to and after 1s stimulation. (F,G) Same as (C,D) but with 25s stimulation. (H) Quantification of categorical neuron responses to optogenetic stimulation series for *Bdnf*^*WT*^ and *Bdnf*^*Nd1-cKO*^ co-cultures. Nonresponders failed to show immediate response to any stimulus. Consistent responders displayed Ca^2+^ spike after every stimulus. Data are mean proportions from 3 *Bdnf*^*WT*^ samples (131 neurons) and 2 *Bdnf*^*Nd1-cKO*^ samples (94 neurons). Scale bars, 250 µm (A, B left panels), 100 µm (A, B middle and right panels), 20 µm (E). *P* values determined by Fisher’s exact test (H). **** *P* < 0.0001.

To determine whether the nodose response scaled with varying levels of EEC activation, we prolonged the period of blue light exposure. In 4-day old *Vglut2*^*GCaMP6s*^ nodose co-cultured with a *Nd1*^*ChR2*^ *Bdnf*^*WT*^ intestine, 50% of measured neurons produced a consistent, evoked calcium spike immediately following a 1s, 5s, and 25s stimulation (Fig. 4F; fig. S8). In contrast, only 15% of nodose neurons cultured alongside *Nd1*^*ChR2*^ *Bdnf*^*Nd1-cKO*^ intestines showed three consistent responses to stimulation (Fig. 4G-H; fig. S8). These “triplet” responders were absent from 1-day old co-cultures, as well as 4-day old “no gut” controls, further suggesting that contact with EECs is required for the evoked calcium spikes. Furthermore, a 25 second stimulation resulted in prolonged activity of nodose neurons relative to the shorter stimulations (Fig. 4F; fig. S7G-I). Nodose neurons cultured alone showed no change in GCaMP fluorescence with prolonged stimulations (fig. S7H,J), while *Nd1*^*ChR2*^ *Bdnf*^*Nd1-cKO*^ co-cultures showed predominantly inconsistent or no response to increased stimulation (fig. S7K). These data confirm that BDNF-dependent, direct interactions with nodose neurons are required for EECs to evoke rapid, consistent, and scalable neuronal activity.

## Direct EEC-nodose communication is essential for neonatal physiology and behavior

Heterozygous mutations in *NTRK2* or haploinsufficiency in *BDNF* have been linked to adolescent obesity, including WAGR syndrome (*11, 12*). To explore how an impaired EEC-nodose circuit impacts physiology, we used *Villin (Vil1*) Cre (*28*) to selectively ablate *Bdnf* in the intestinal epithelia and monitored neonatal weight gain. Our scRNAseq analyses of intestinal epithelia revealed that only EECs express *Bdnf* (Fig. 2C), meaning this *Vil1*^*Cre*^; *Bdnf*^*fl/fl*^ (*Bdnf*^*Vil1-cKO*^) model serves as an EEC-specific *Bdnf* deletion without affecting any non-intestinal tissues. Despite no significant difference in birthweight, neonatal *Bdnf*^*Vil1-cKO*^ mice were consistently underweight relative to their *Bdnf*^*WT*^ littermates from P1-P15, when mice exclusively suckle (Fig. 5A). These data suggest that formation of the EEC-nodose circuit, *in utero*, is essential to early weight gain—possibly by regulating suckling behavior.

**Figure 5.**
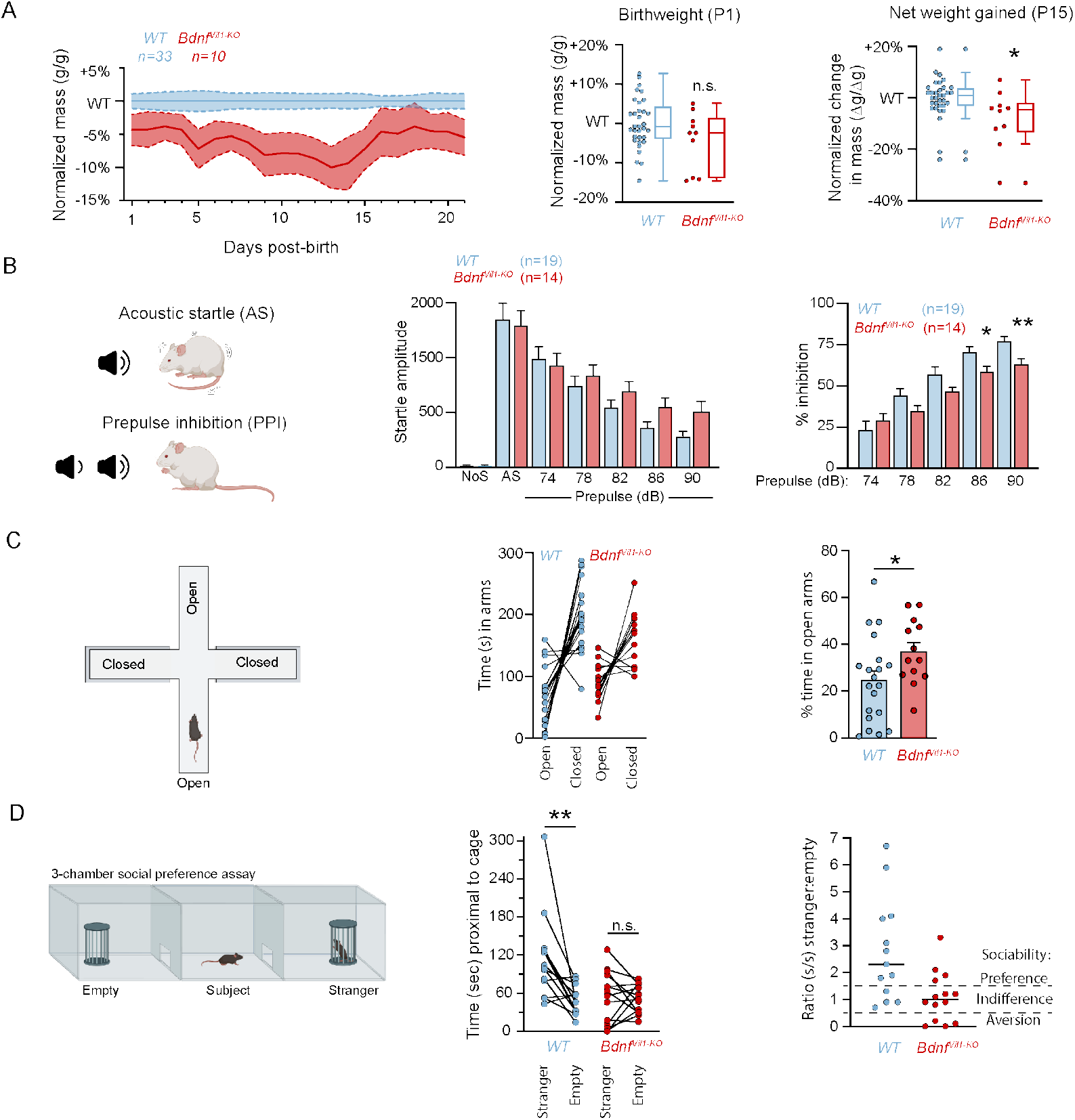
Deficits in weight, sociability, and inhibitory control upon intestine-specific *Bdnf* knockout. (A) Neonatal weight gain for *Bdnf*^*Vil1-cKO*^ individuals (red) and WT littermates (blue). Left chart displays daily neonatal mass from P1-P21 (birth through weaning) normalized to average mass of wild-type individuals within each litter; *n* refers to the number of neonates from 4 separate litters. Middle graph shows birthweight normalized to the average P1 mass of wild-type individuals within each litter. Dots represent individual neonates. Right chart displays quantification of mass gained by P15 (end of suckling exclusivity) normalized to the average mass gained of wild-type individuals within each litter. (B) Acoustic startle response assay depicting a schematic of the acoustic chamber (left), startle amplitude (middle), and % inhibition to the prepulse (right). Prepulse of increasing dBs increasingly reduces the startle reflex in response to AS (120 dB). *Bdnf*^*Vil1-cKO*^ individuals display reduced prepulse inhibition, suggesting deficits in sensorimotor gating. NoS = no sound. AS = acoustic stimulus (120 dB). (C) Elevated plus maze (EPM) assay. (Left) Schematic of EPM apparatus. (Middle) Paired measurements of time (in seconds) spent in the open and closed arms of the elevated plus maze. (Right) Percent of time spent in the open arms of the elevated plus maze. *Bdnf*^*Vil1-cKO*^ individuals display increased risky (open arm) exploration. (D) 3-chamber social preference assay used to assess sociability, showing a schematic (left), and measurements of time spent (middle) and ratios (right) within 5 cm proximity to stranger vs empty cages. Most WT individuals displayed social preference (>50% increase in time spent near stranger cage) while most *Bdnf*^*Vil1-cKO*^ individuals displayed social indifference (stranger:empty ratio = 1±0.5) or social aversion (>50% decrease in time spent near stranger). * *P* < 0.05, ** *P* < 0.01 determined by student’s unpaired t-test (A-C) or Fisher’s PLSD test (D).

*BDNF* and *NTRK2* are implicated in the pathophysiology and treatment of a variety of neurological disorders **(*22*)**. Mice undergoing a subdiaphragmatic vagal deafferentation, which selectively severs the afferents projecting to the abdominal viscera, display hallmarks of schizophrenia including a loss of prepulse inhibition (PPI) in an acoustic startle test (*29*). *Bdnf*^*Vil1-cKO*^ animals exhibited reduced PPI compared to *Bdnf*^*WT*^ littermate controls, indicative of deficits in sensorimotor gating (Fig. 5B), further supporting that *Bdnf* deletion selectively impairs EEC interactions with vagal afferent fibers.

Because BDNF and gut-brain communication have been implicated in the development of anxiety and stress responses (*30, 31*), we assessed *Bdnf*^*Vil1-cKO*^ animals for altered anxiety-like behaviors in an elevated plus maze. *Bdnf*^*Vil1-cKO*^ animals spent a larger percentage of their time in the open arms compared to *Bdnf*^*WT*^ littermates (Fig. 5C) and entered the open arms more frequently (fig. S9A,B), suggesting that the EEC-nodose circuit regulates anxiety-like responses. This behavior could not be explained by general hyperactivity, because *Bdnf*^*Vil1-cKO*^ littermates showed no difference in total arm entries (fig. S9C). The observed variance in both PPI and the elevated plus maze suggest that compromised EEC-nodose communication may lead to generalized deficits in inhibitory behavioral control.

Circulating levels of BDNF correlate with the development of autism spectrum disorders (ASD) (*13*), which are characterized by a lack of sociability and repetitive behaviors. Sociability defects in mouse models of ASD can be rescued by altering the microbiome, and this effect requires an intact vagus nerve (*32*). ASD patients also present with alterations in their gut microbiome and frequently display gastrointestinal comorbidities, yet whether GI-specific genetic alterations can contribute to ASD symptoms is unknown (*33*). To test this, we evaluated *Bdnf*^*Vil1-cKO*^ mice for sociability using a 3-chamber social preference assay. While *Bdnf*^*WT*^ animals consistently interacted with a stranger mouse more than an empty cage, their *Bdnf*^*Vil1-cKO*^ littermates often spent equal time exploring both cages, or more time investigating the empty cage (Fig. 5D). These differences are not due to lack of exploratory behavior, visual deficits, hyperactivity, or locomotor ability as *Bdnf*^*Vil1-cKO*^ littermates showed no variance in total side entries (fig. S9D,E), performance in a visible platform test (fig. S9F), and total distance traveled or rearing movements (fig. S9G,H).These data mirror established mouse models of ASD and suggest that proper formation of the EEC-nodose circuitry is essential to the development of normal sociability. Collectively, these studies are the first to suggest that impaired development of gut-brain circuits may contribute not only to feeding behaviors, but also to symptoms of anxiety, autism and other neurodevelopmental disorders.

## DISCUSSION

Our studies define the ontogenesis of mammalian gut-brain communication and uncover novel behaviors regulated by the EEC-nodose circuit. That *Bdnf* plays a key role in regulating gut-brain communication via the interaction between EECs and vagal nodose neurons has far reaching implications for both the development and treatment of a variety of gut-brain disorders, including obesity and ASD. The fact that *Bdnf* ablation specifically in the intestinal epithelia is sufficient to alter not only weight gain, but also animal behavior, highlights the importance the gut— and direct EEC-nodose communication in particular—in affecting neurological phenotypes.

In adult mice, optogenetic or chemogenetic activation of CCK+ EECs suppresses feeding behavior (*1, 9*). In humans, heterozygous mutations in *BDNF* or *NTRK2* cause obesity (*11, 12*), as does haploinsufficiency of *Bdnf* in mice (*34*). In contrast, our finding that *Bdnf*^*Vil1-cKO*^ neonates are underweight suggests that the interaction of *Bdnf*-expressing EECs and nodose neurons promotes, rather than inhibits, suckling (Fig. 5A-C). While this result is counterintuitive, it should be noted that the role of a circuit can change across development. For example, hypothalamic Agrp neurons are essential for feeding in adults (*35*), but dispensable during early development where they instead regulate maternal association (*36*). Moreover, prior studies modulated EEC activity *en masse*, suggesting that contact-based EEC-nodose communication may have unique effects on behavior. Lastly, the observation that our *Bdnf*^*Vil1-cKO*^ mice were underweight while patients with heterozygous mutations are obese highlights that different effects may be observed upon loss versus reduction of this pathway.

The links between gastrointestinal and neurological disorders are too numerous to detail here and have been extensively reviewed elsewhere (*37, 38*). However, it is worth noting that there is increased interest in the role of the vagus nerve, given its place as a conduit between these two systems. Studies have shown the intestinal microbiome can play a causal role in ASD symptoms (*32, 39*), and our results suggest that these effects may be mediated by *Bdnf*-dependent EEC-nodose communication. Furthermore, BDNF and its receptor, TrkB, have wide ranging implications in both the development and treatment of neurological disorders, including depression, anxiety, and ASD (*13, 22*). In addition, it has recently been shown that anti-depressants can not only inhibit serotonin reuptake, but also activate TrkB as allosteric potentiators (*40*). Taken together, these findings place BDNF-TrkB and the EEC-nodose circuit at the nexus of behavior and gut-brain pathophysiology.

## Acknowledgments

We thank members of the Williams lab, Jose Rodriguez-Romaguera, Pablo Ariel, John Rawls, Stephanie Gupton, Laura Rupprecht, and Kaelyn Sumigray for their critical feedback. We thank Dr. Pablo Ariel for his support performing and analyzing the optogenetic experiments. We thank Dr. Kathryn Harper and Samuel Harp M.S. for their support in performing the behavioral assays. We thank Dr. Diego Bohórquez for graciously sharing rabies viral stocks for early pilot studies of our *in utero* transsynaptic tracing. We thank Dr. Melanie Kaelberer for her guidance performing nodose ganglia dissections and neonatal oral gavage. We thank Dr. Zachary Knight for sharing the *Bdnf-2A-Cre* mouse strain with us. We thank Dr. David Ginty for sharing the TrkB-flox and TrkB-tauGFP mouse strains with us.

KJL was supported by the UNC Center for Gastrointestinal Biology and Disease (CGIBD) T32 Basic Science GI training fellowship (National Institutes of Health grant T32 DK07737) and a UNC CGIBD pilot feasibility award (supported by National Institutes of Health grant P30 DK034987). SW is supported by an R01 from the National Institutes of Health, National Institute of Arthritis and Musculoskeletal and Skin Diseases (R01 AR077591).

The Andor Dragonfly microscope was funded with support from National Institutes of Health grant S10 OD030223. The single cell RNA sequencing and FACS isolation were performed in the UNC CGIBD Advanced Analytics Core, using equipment supported by NIH grant P30 DK034987. The behavioral studies were performed in facilities supported by the Eunice Kennedy Shriver National Institute of Child Health and Human Development grant P50 HD103573. The rabies viruses were produced by the UNC NeuroTools viral vector core facility, which receives support from the NIH BRAIN Initiative (U24 NS124025-04).

## Data and materials availability

Raw and processed scRNAseq data have been deposited with the Gene Expression Omnibus (accession number GSE283137). This dataset is currently private, but data will be made public upon acceptance of this manuscript for publication.

## Supplementary Materials

Materials and Methods

Figs. S1 to S9

Table S1

Movies S1 to S4

## Supplementary Materials

### Materials and Methods

Table S1 contains a list of all reagents, including vendor information, catalogue and RRID numbers.

#### Mice

Mice were housed in an AAALAC-accredited (#329; November 2023), USDA registered (55-R-0004), NIH welfare-assured (D16-00256 (A3410-01)) animal facility. Mice were housed in standard housing under 12-12 hour light-dark cycles with *ad libitum* access to standard chow and water with access to enrichment. Only male breeders were singly-housed, all other mice were co-housed with 2-5 adult littermates per cage. All lines used were maintained on a mixed background consisting of C57BL/6J (Jackson) and CD1 (Charles River) lab strains. All experiments were performed using littermate controls, with at least two litters. All studies employed a mixture of male and female mice, though this was not explicitly tracked. Pregnancies were timed by both plug-checking and ultrasound imaging using a ViewSonics Vevo 2100. Note that *Neurod1*^*Cre*^ (Jax #: 028863) is a knock-in, loss-of-function allele, so these mice were maintained as heterozygotes and never bred to another carrier of the allele. Other Cre alleles are fully functional. The *Bdnf*^*2A-Cre*^ line (*52*) was a generous gift from Zachary Knight, all other strains were procured from Jackson labs.

#### Single-cell RNA dataset

Embryonic EECs were isolated from E15 (3 embryos), E16 (4 embryos) and E17 (4 embryos) *Neurod1*^*mGFP*^ embryos. Briefly, intestines were dissected out and bisected longitudinally to expose the luminal epithelia and submerged in cold dissociation cocktail (2.25 mg/mL DNAse (Sigma Cat. #: DN25), 10 mg/mL protease (Sigma Cat. #: P5380), and 9 µM Y-27632 (Selleck Chem. Cat. #: S1049) in PBS). Samples were incubated in dissociation cocktail on ice for 30 minutes with trituration every 5 minutes until achieving >90% single cells in the suspension. Cells were then rinsed 3x in cold PBS, passed through a 40 µm cell strainer, pelleted and resuspended in cold DMEM+10% FBS with 2.25% DNAse and 9 µM Y-27632 to preserve viability during cell sorting. Final cell counts were ∼2.75e6 cells from E15.5 embryos, ∼7.5e6 cells from E16.5 embryos and ∼7.5e6 cells from E17.5 embryos, all of which were stained with Sytox and Annexin V (to label dead cells) and pooled prior to sorting. FACS isolation of GFP^POS^ EECs was performed on a Sony SH800 cell sorter with gating based on GFP^NEG^ controls (one *Cre*^*NEG*^ *ROSA*^*mTmG*^ littermate from each age), excluding Sytox^POS^ and Annexin V^POS^ cells. Sorted ∼65,000 GFP^POS^ cells and viability was determined to be ∼89% by LUNA-FL Cell Counter before library preparation.

10,000 single cells were processed through the Chromium Next GEM Single Cell 3’ Reagent Kit v3.1 (Dual Index) with Feature Barcode technology for Cell Surface Protein (10X Genomics, Pleasanton, CA) as per the manufacturer’s instructions. Single-cell cDNA libraries were sequenced on the Illumina NextSeq 2000 platform with a NextSeq P3 flow cell. De-multiplexing alignment to the mm10-2020-A transcriptome and unique molecular identified (UMI)-collapsing were performed using Cell Ranger (v7.0.0). A gene-barcode matrix was generated with ∼5.77e8 reads over 8,912 cells (∼65,000 mean reads per cell and ∼3,000 mean genes detected per cell).

#### Single-cell RNA analysis

Pre-processing and data visualization were performed using Seurat (v3), following standard workflows. We excluded cells with >5% mitochondrial reads, fewer than 1,000 RNA counts, or fewer than 500 genes detected, resulting in 6,061 total cells. The filtered dataset was normalized using the SCTransform function and clustered with a Louvain resolution of 0.5. Louvain resolution was determined based on the ability to recreate previously reported subpopulations of EECs (*9, 16*). Cluster identification was based on published marker genes including enterochromaffin cells (*Tac1*^*+*^, *Tph1*^*+*^), I cells (*Cck*^*Hi*^), K cells (*Cck*^*+*^, *Gip*^*Hi*^), N/L cells (*Cck*^*+*^, *Pyy*^*Hi*^, *Gcg*^*Hi*^), X cells (*Ghrl*^*+*^, *Sst*^*NEG*^), D cells (*Ghrl*^*+*^, *Sst*^*+*^), neurons (*Ncam1*^*+*^, *Snap25*^*+*^), enteric glia (*Nfia*^*+*^), epithelia (*Epcam*^*+*^, *Vil1*^*+*^, *Cdh1*^*+*^), and enterocytes (*Fabp1*^*+*^).

#### *in utero* rabies injections

The protocol used in this study was approved via IACUC #22-121. Breeder pairs were set up with one *Nd1*^*Cre/+*^; *ROSA*^*RABvG/RABvG*^ parent bred to a *ROSA*^*mTmG/mTmG*^ partner. Pregnancies were timed by ultrasound monitoring and measuring crown-rump length. Pregnant dams carrying E13.5 or E15.5 embryos were anesthetized and the uterine horn pulled into a PBS filled dish. Embryos and custom glass needles were visualized by ultrasound (Vevo 2100) to guide microinjection of ∼1.0 μl of concentrated rabies viral solution into the umbilical hernia. The viral solution was diluted to ∼1e7 infectious units per μl, with 1 μl of red fluorescent beads (FluoSpheres; ThermoFisher Cat. #: F8801) per 10 μl of viral solution. Prior to addition, fluorescent beads were rinsed 3x in sterile PBS to remove sodium azide. Three to eight embryos were injected depending on viability and litter size. Following injection, the uterine horn(s) were reinserted into the mother’s thoracic cavity, which was sutured closed. The incision in the skin was resealed with surgical staples and the mother provided subcutaneous analgesics (5 mg/kg meloxicam and 1-4 mg/kg bupivacaine). Once awake and freely moving, the mother was returned to its housing facility for 3-5 days, at which point E16.5-18.5 embryos were harvested and processed accordingly.

#### Live Imaging

A 1% agar solution/media solution containing F^+^-media (3:1 DMEM:F12 + 10% FBS + 1% Sodium bicarbonate + 1% Sodium Pyruvate + 1% Pen/Strep/L-glut mix, + 1% B-27 and 2% N2 neurogenic supplements), was cooled and cut into 35 mm discs. Proximal intestine samples (∼1-2 mm in length) were dissected from E15.5 embryos and cut longitudinally to expose the gut lumen. These were placed lumen-side up onto the gel/media disc, then sandwiched between the gas-permeable membrane of a 35 mm lumox culture dish (Sardstedt Cat. #: 94.6077.331). For gut-ganglion cocultures, the nodose ganglion was dissected from an E15.5 embryo and the nerve fibers were trimmed as short as possible while avoiding damaging the ganglion. The ganglion was placed directly adjacent to the gut tissue. Live imaging was performed utilizing an Andor Dragonfly laser spinning disk confocal microscope equipped with a Zyla Plus 4.2 MP cMOS camera. For innervation and EEC-neuron contact studies, images were acquired with 5-minute intervals, with a voxel size of 0.314 µm x 0.314 µm x 0.494 µm, and a z-stack that included all observable EECs with Nyquist sampling (0.297 µm spacing). Samples were imaged daily for 7 days for 2-4 hours using an HC PL APO 20x/0.75 LWD air objective. For optogenetic-GCaMP6s studies, images were acquired with 500 ms intervals in a single plane using an HC PL APO 10x/0.45 air objective with a pixel size of 0.628 µm x 0.628 µm, in a single z plane. mGFP, ChR2-YFP and GCaMP6s were excited using 488 nm laser (3% laser power, 450 ms exposure, with the F521-038 emission filter), while mTdTomato was excited with a 561 nm laser (4% power, 450 ms exposure, with the F594-043 emission filter). Optogenetic stimulation was achieved using 405 and 470 nm LEDs (CoolLED pE-4000), regionally restricted to the intestinal sample using a MOSAIC digital mirror device (Oxford Instruments). Samples were serially imaged as follows: **Protocol 1:** 10 minutes of pre-stimulation, 1s blue-light stim, 10 minutes of post-stimulation. **Protocol 2:** 2 minutes of pre-stimulation, 5s blue-light stim, 10 minutes of post-stimulation. **Protocol 3:** 2 minutes of pre-stimulation, 25s blue-light stim, 10 minutes of post-stimulation. Explants were cultured at 37°C with 5.0% CO_2_ throughout the course of the experiment(s).

#### Antibodies, immunohistochemistry, and fixed wholemount imaging

Intestines and nodose ganglia were dissected from E15-E18 embryos and fixed in 4% room temperature paraformaldehyde overnight at 4C, then washed 3×30 minutes with PBS + 0.4% Triton X-100 (PBSTx) and stored in PBS + 0.05% sodium azide at 4°C. Intestinal samples were bisected longitudinally before fixation. Prior to staining, proximal, mid, or distal gut samples were dissected into smaller, 1-5mm pieces and submerged overnight at 4°C in wholemount gelatin block (5% NDS, 3% BSA, 8% cold-water fish gelatin, 0.4% Triton X-100 in PBS). Primary antibodies were diluted in wholemount gelatin block and samples were submerged in primary antibody solution for 36-72 hours at 4°C. Samples were then washed with PBSTx and incubated in secondary antibodies diluted in wholemount gelatin block overnight at 4°C, then counterstained with DAPI (1:2000) overnight at 4°C and mounted in ProLong Gold (Invitrogen). Images were acquired using LAS AF software on a Leica TCS SPE-II 4 laser confocal system on a DM5500 microscope with ACS Apochromat 20x/0.60 multi-immersion, or ACS Apochromat 40x/1.15 oil objectives. All images for assessing EEC-neuron contact were taken with the 40x/1.15 oil objective and a voxel size of 0.269 µm x 0.269 µm x µm x 0.5 µm, therefore, instances of “contact” are defined as <500 nm distance between EEC mGFP+ membrane and Tuj1 stained axons, and often represent closer approximations (e.g. if observed in x,y plane).

The following primary antibodies were used: monoclonal anti-mCherry (rat, ThermoFisher M11217, 1:1000-3000), polyclonal anti-GFP (chicken, Abcam ab13970, 1:1,000), monoclonal anti-Tuj1 (mouse, Abcam ab78078, 1:500), polyclonal anti-CCK (rabbit, Phoenix Pharmaceuticals G-069-04, 1:1,000), monoclonal anti-SubP (rat, Millipore MAB356, 1:250), polyclonal anti-TrkA (rabbit, Millipore 06-574, 1:300), polyclonal anti-TrkB (goat, R&D Systems AF1494, 1:500), polyclonal anti-Phox2b (goat, R&D Systems AF4940, 1:500).

The following secondary antibodies were used (all antibodies produced in donkey): anti-rabbit AlexaFluor 488 (Life Technologies, 1:1000), anti-rabbit Rhodamine Red-X (Jackson Labs, 1:500), anti-rabbit Cy5 (Jackson Labs, 1:400), anti-rat AlexaFluor 488 (Life Technologies, 1:1000), anti-rat Rhodamine Red-X (Jackson Labs, 1:500), anti-rat Cy5 (Jackson Labs, 1:400), anti-goat AlexaFluor 488 (Life Technologies, 1:1000), anti-goat Cy5 (Jackson Labs, 1:400), anti-mouse IgG AlexaFluor 488 (Life Technologies, 1:1000), anti-mouse IgG Rhodamine Red-X (Jackson Labs, 1:500), anti-mouse IgG Cy5 (Jackson Labs, 1:400), anti-Chicken AlexaFluor 488 (Life Technologies, 1:1000).

#### Behavior assays

Mice were subjected to a battery of behavioral assessments in the order listed below. More stressful procedures were delayed toward the end of the series to avoid confounding effects on other assessments. Mice were evaluated in no more than two different assays per week. Assessments were performed across three separate cohorts of mice, with all mice being group-housed with 3-4 littermates per cage, and each cage contained some number of both wild-type and *Bdnf*^*Vil1-cKO*^ animals. Testing started when mice were 6-9 weeks in age. Subjects were 19 wild-type individuals (9 female, 10 male) and 14 *Bdnf*^*Vil1-cKO*^ animals (6 female, 8 male).

#### Elevated plus maze

Mice are place on an elevated (50 cm from floor) maze with a central chamber (8 × 8 cm) and four arms (each 30 cm long). Two of the arms are closed (20 cm high walls) and two of the arms are open (no walls). During a 5-mintue trial, mice were allowed to freely explore the maze and measures were taken of 1) the number of entries into, and 2) time spent in each arm. *Open field assay*. Mice were placed in an open chamber (41 × 41 × 30cm) crossed by a grid of photobeams to report the animal’s position within the chamber (VersaMax, AccuScan Instruments). During a 1-hour trial, the counts of all photobeams broken are assessed to evaluate 1) the total distance traveled, 2) number of rearing movements, and 3) amount of time spent in the center of the chamber versus 4) amount of time spent near the chamber walls.

#### 3-chamber social preference assay

Mice were placed in a rectangular social testing apparatus consisting of three-chambers constructed of clear Plexiglas, with doorways allowing access into each chamber. Each mouse was granted a 10-minute habituation period where they were allowed to explore the empty social test box with all doors open. Next, the mouse is confined to the center chamber, and 2 Plexiglas cages drilled with small holes are placed in each of the side chambers. One cage is empty, and the other cage contains an unfamiliar, sex-matched mouse (stranger). The doors for the center chamber are then re-opened, and the test animal is allowed to freely explore the social test box for 10 min. An automated image tracking system (Noldus Ethovision) provided measures of 1) entries into each side chamber, 2) time spent in each side chamber, and 3) time spent in close (<5cm) proximity to each cage. Data were removed for one mouse which never left the central chamber, and six additional mice which did not explore the wire cages (<20 sec total time spent proximal to cages; removed data were from 5 wild-type mice, and 1 heterozygote [data not shown]). Inclusion of these individuals adjusts the p value for *Bdnf*^*WT*^ time spent in proximity to stranger vs empty cage from p=0.0072 to p=0.0138 (i.e. no change in significance).

#### Acoustic startle and prepulse inhibition

Mice were placed in the SR-LAB-Startle Response System (San Diego Instruments), which consists of a small Plexiglas cylinder housed within a larger noise-dampening chamber. The Plexiglas cylinder rests upon a piezoelectric transducer which records change in force applied during the whole-body reflex. Subjects were allowed a 5-minute habituation period, followed by 6-replicates of 7 trials: no-stimulus (NoS), a 40 msec, 120 dB acoustic startle stimulus alone (AS), and prepulse trials where a 20 msec acoustic pulse of varying magnitude (74, 78, 82, 86, or 90 dB), precedes the acoustic startle by 100 msec. The trial types are randomized within each trial series, with an average intertrial interval of 15 seconds. The startle response is measured as the peak startle amplitude during a 65 msec window coincident with the onset of the startle stimulus.

#### Morris water maze

The water maze was used to assess swimming ability and vision. The water maze consisted of a large circular pool (diameter = 122 cm) partially filled with water (45 cm deep, 24-26°C), located in a room with numerous visual cues. In the visible platform test, each mouse was given 4 trials per day, across 2 days, to swim to an escape platform cued by a patterned cylinder extending above the surface of the water. For each trial, the mouse was placed in the pool at 1 of 4 possible locations (randomly ordered), and then given 60 sec to find the visible platform. If the mouse found the platform, the trial ended, and the animal was allowed to remain 10 sec on the platform before the next trial began. If the platform was not found, the mouse was placed on the platform for 10 sec, and then given the next trial. Measures were taken of latency to find the platform and swimming speed via an automated tracking system (Noldus Ethovision).

#### Measurements, quantification, graphing, and statistics

##### EEC density and subtype prevalence

To determine EEC density, we first counted individual EECs (GFP+ epithelia from *Nd1*^*mGFP*^ samples) in 3D, volumetric images from wholemount samples using the ImageJ CellCounter plug-in. We separately counted CCK+ and SubP+ epithelia. Importantly, all imaged E15-E17 proximal, mid, and distal samples were treated with the same antibody solutions prepared on the same day. To determine sample volume, we generated a 3D surface of the DAPI signal in Imaris, using a minimal threshold to generate the surface in a manner reflecting the tissue shape. To avoid differences in imaging volume confounding results, we only imaged the full villus thickness of the tissue and avoided imaging the submucosa. We then divided the counted EEC populations by the volume of the tissue, standardizing the units to # of EECs (of a given type) per (100 µm)^3^ (equivalent to a cube where each side is 100 µm; i.e. 1e6 µm^3^). For reference, the max-projection images in Fig. 1C are 2D projections equivalent to 2e6 µm^3^ or 2 units of (100 µm)^3^. The proportion of EEC subtypes across ages and intestinal regions were quantified by dividing counted CCK+ or SubP+ epithelia by total mGFP+ epithelia from *Nd1*^*mGFP*^ embryos (see Fig. 1F).

##### EEC-neuron contact frequency

Determination of “contact” between EECs and neurons was dependent on observation of overlapping pixels between mGFP and Tuj1 in samples taken from *Nd1*^*mGFP*^ embryos. To avoid detection of false positives and control for background signal, Tuj1 fluorescence was thresholded to the point of observing discrete neural fibers. We then counted the total number of observed instances of contact per villus for 4 individual embryos at E15 and E16, plotting the mean number for each individual in Fig. 1J. The proportion of EECs with contacts was determined by counting all EECs with contact and dividing that population by the total number of EECs for the same four individuals. Contacts per villus and proportion of EECs with Tuj1 contact were compared by unpaired student’s t-test.

##### GCaMP6s fluorescence

ROIs were hand drawn around any soma with observable calcium flares during the imaging session and the mean fluorescence intensity of all pixels within the ROI throughout the extent of the experiment was measured. To correct for sample drift and photobleaching, we measured baseline GCaMP6s fluorescence (F_0_) in a rolling 2-minute window as the average of the lowest 20 mean fluorescence values in a given ROI and subtracted this value from the raw fluorescence mean within the ROI at every timepoint (F-F_0_= ΔF). We corrected for bleed through of the stimulation light as outlined in Fig. S6C. Briefly, for the first 1 sec at stimulation onset (two frames), we set F_0_ as the mean intensity in the ROI at the exact onset of the stimulation (F*; hence, the stimulation frame always has ΔF/ΔF_MAX_=0, which is why the stimulation frame is grayed out in Fig. 6E, G, H). For the 5 sec stimulation, we again set F_0_ as F* for the first two frames. For the remaining 4 seconds while the blue light was ON, we re-calculated F_0_ as the mean of the lowest 5 values detected during the remainder of the 5 sec stimulation. For the 25 sec stimulation, we again set F_0_ as F* for the first two frames. For the remaining 24 seconds while the blue light was ON, we re-calculated F_0_ as the mean of the lowest 10 values detected during the remainder of the 25 second stimulation. These values were normalized to the maximum value of every cell (scale 0-1) by dividing each ΔF by the max ΔF (ΔF/ΔF_MAX_). ΔF_MAX_ was the maximum in the entire experiment (including all 10 min rests and 2 min stimulation periods; total 36 minutes), and we did not dictate whether ΔF_MAX_ was either a spontaneous or evoked spike. Immediate responders were counted as cells that displayed a value of at least 0.25 ΔF/ΔF_MAX_ within 1 second of the start of stimulation. To compare the scalability of responses during prolonged stimulation, we summed the ΔF/ΔF_MAX_ for the 30 seconds after the onset of blue light stimulation.

##### Neonatal weight

Mice were weighed daily from birth (day litter was first observed, assumed to be P1) until weaning (P21). Heterozygous breeder pairs produced litters containing at least four WT individuals and at least one *Bdnf*^*Vil1-cKO*^. Data presented are from six litters. All measurements were normalized to the average for WT individuals within the litter. Weight gained is the difference in mass between P15 and P1, normalized to the average mass gained for WT littermates. Comparisons of weight gained and birthweight were tested by unpaired student’s t-test. We found no significant difference in weights between sexes until after P21.

##### Acoustic startle and PPI

The startle amplitude was measured as the peak startle amplitude during a 65 msec window coincident with the onset of the startle stimulus. Prepulse inhibition (PPI) was measured by dividing the startle amplitude with the pre-pulpse by the startle amplitude for the acoustic startle (120 dB) alone. This proportion was subtracted from 1, then multiplied by 100 to generate the % prepulse inhibition [100*(1-(prepulse startle/AS))]. The *Bdnf*^*Vil1-cKO*^ mice had significant reductions in prepulse inhibition in comparison to *Bdnf*^*WT*^ controls [genotype x decibel level interaction, F(4,124)=3.84, p=0.0056].

##### Elevated plus maze

Percent time in open arms was quantified by dividing the time (in seconds) spent in the open arms by the total time spent in either arms [100*(open time/(open time+closed time))]. In this manner, time spent in the central chamber was excluded from the quantification. The *Bdnf*^*Vil1-cKO*^ mice had higher percent open arm time [F(1,31)=4.25, p=0.0477] and percent open arm entries [F(1,31)=6.6, p=0.0152] in the elevated plus maze, compared to *Bdnf*^*WT*^.

##### Sociability

Stranger to empty cage ratios were calculated by dividing the time (in seconds) the subject spent proximal (<5 cm) to the stranger cage by the time spent proximal to the empty cage. The *Bdnf*^*WT*^ mice spent significantly more time in proximity to the stranger cage vs empty cage [within genotype, F(1,12)=10.43, p=0.0072] while *Bdnf*^*Vil1-cKO*^ mice did not have positive sociability [within genotype, F(1,13)=0.03117, p=0.8626].

All statistical analyses and graphs were generated using GraphPad Prism. Error bars represent standard error of the mean (s.e.m.) unless otherwise noted. Statistical tests of significance were determined by Mann-Whitney *U*-test (non-parametric) or student’s t-test (parametric) depending on whether the data fit a standard distribution (determined by pass/fail of majority of the following: Anderson-Darling, D’Agostino & Pearson, Shapiro-Wilk, and Kolmogorov-Smirnov tests). χ^2^ tests were utilized to evaluate expected (control) against experimental distributions of categorical values (e.g. EEC-nodose interaction behaviors in explant cocultures). For all behavior assays, data were first analyzed using two-way or repeated measures Analysis of Variance (ANOVA), with the factors genotype and sex. Fisher’s protected least significant difference (PLSD) tests were used for comparing group means only when a significant F value was determined (we did not observe significant F-values for sex). All box-and-whisker plots are displayed as Tukey plots where the box represents the interquartile range (IQR, 25^th^-75^th^ percentiles) and the horizontal line represents the median. Whiskers represent 1.5x IQR unless this is greater than the min or max value. Figures were assembled using Adobe Photoshop and Illustrator CC 2024-2025.

**Fig. S1.**
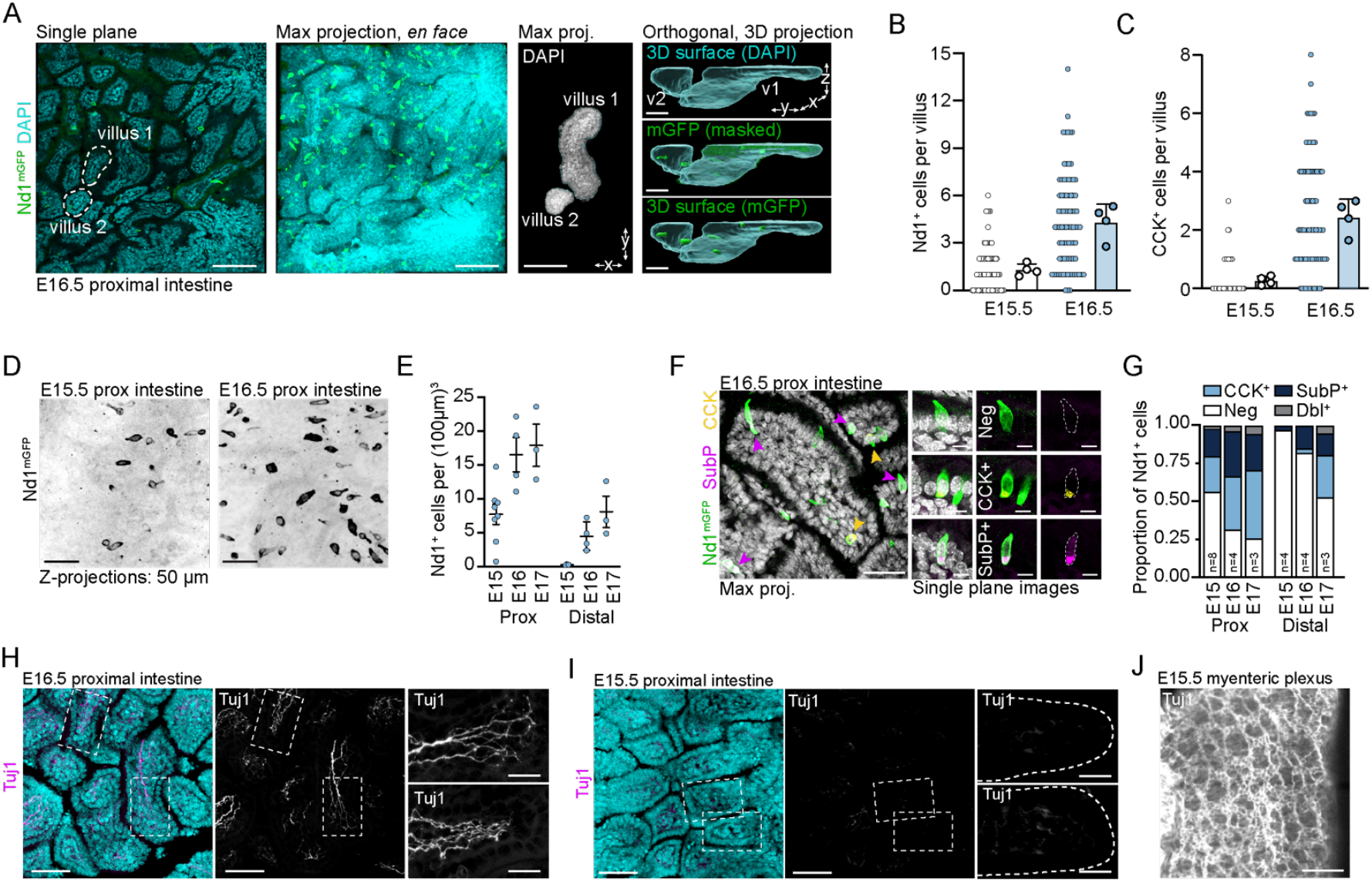
Development of embryonic EECs and early myenteric innervation. (A) Wholemount fluorescence microscopy of the embryonic day 16.5 (E16.5) intestine. Left to right: single focal plane of proximal intestine; max z-projection of the same field; DAPI max pojection of isolated villi for dotted region in leftmost image; Imaris based 3D projections of villi. (B) Quantification of the frequency of Nd1+ cells per villus from E15.5 and E16.5 proximal intestines. Small dots at left represent single villus; large dots at right represent average per embryo. Data presented are from 4 individuals and at least 2 separate litters. (C) Quantification of the frequency of CCK+ cells per villus from E15.5 and E16.5 proximal intestines, shown as in (B). (D) Max projections of mGFP (black) proximal intestines from *Nd1*^*mGFP*^ embryos at E15.5 (left) or E16.5 (right). (E) Nd1+ cell (EEC) density based on embryonic age and intestinal region. Each dot represent data from one individual embryo, with at least 2 separate litters represented per timepoint. (F) Wholemount E16.5 proximal intestine stained for *Nd1*^*mGFP*^ (green), substance P (SubP; magenta) and cholecystokinin (CCK; yellow) showing max projection (left) and single plane images of individual EECs (right). (G) Quantification of EEC subtype based on accumulation of CCK and/or SubP within *Nd1*^*mGFP*+^ epithelia; *n* indicates individual embryos from at least two separate litters per timepoint. (H,I) Max projection images of proximal intestine at E16.5 (H) or E15.5 (I) stained for Tuj1. Right-most panels depict individual villi outlined in white dashed boxed in the first two panels. (J) Max z-projection image of E15.5 myenteric plexus from proximal intestine stained for Tuj1 (magenta) to label all axons. Scale bars, 100 m (A, first three panels), 25 µm (A, right), 40 µm (D,F), 75 µm (H-J).

**Fig. S2.**
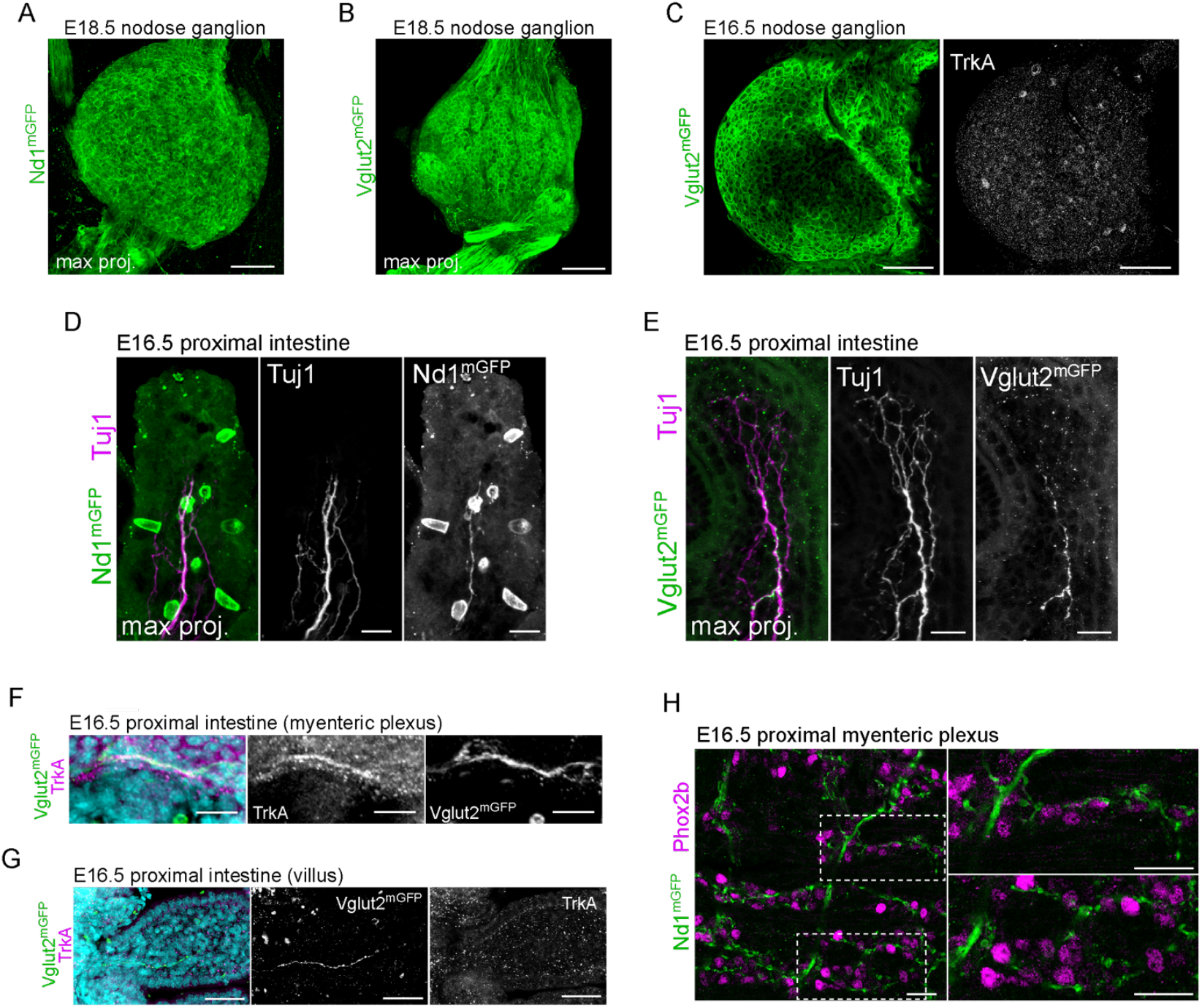
Nodose neurons are the source of extrinsic projections in the embryonic proximal intestinal mucosa. (A-D) Max projection images of wholemount E18.5 nodose ganglia (A,C) and intestinal villi (B,D) from *Nd1*^*mGFP*^ (A,B) or *Vglut2*^*mGFP*^ embryos (C,D). stained for mGFP (green) and Tuj1 (magenta) to label axons. With both markers, mGFP+/Tuj1+ projections can be observed in the intestinal mucosa. (E) Max z-projection image of E16.5 myenteric plexus from proximal intestine of *Nd1*^*mGFP*^ embryo stained with Phox2b (magenta) to label nuclei of enteric neurons and mGFP (green). While mGFP+ projections can be seen in the myenteric plexus, enteric nuclei are not mGFP+ enclosed, suggesting that E16.5 enteric neurons do not express *Neurod1*. (F) Max z-projection image of wholemount E16.5 nodose ganglion from *Vglut2*^*mGFP*^ embryo stained for mGFP (green) and TrkA (gray, right panel). Very few neurons are TrkA+ in the nodose ganglion. (G) Max projection images of E16.5 proximal intestine from *Vglut2*^*mGFP*^ embryo stained for mGFP (green; gray bottom right panel) and TrkA (magenta; gray middle right panel). Right panels are magnification of gut wall region showing mGFP+ projection co-stained with TrkA. (H) Max projection images of E16.5 proximal intestine from *Vglut2*^*mGFP*^ embryo stained for mGFP (green; gray bottom left panel) and TrkA (magenta; gray bottom right panel) showing mGFP+ projection in the villi mucosa. Mucosal endings of mGFP+ projections from *Vglut2*^*mGFP*^ samples do not colabel with TrkA, suggesting they are not DRG projections. Scale bars, 100 µm (A, C, E, F), 20 µm (B, D, E, G right panels), 50 µm (G left panel, H).

**Fig. S3.**
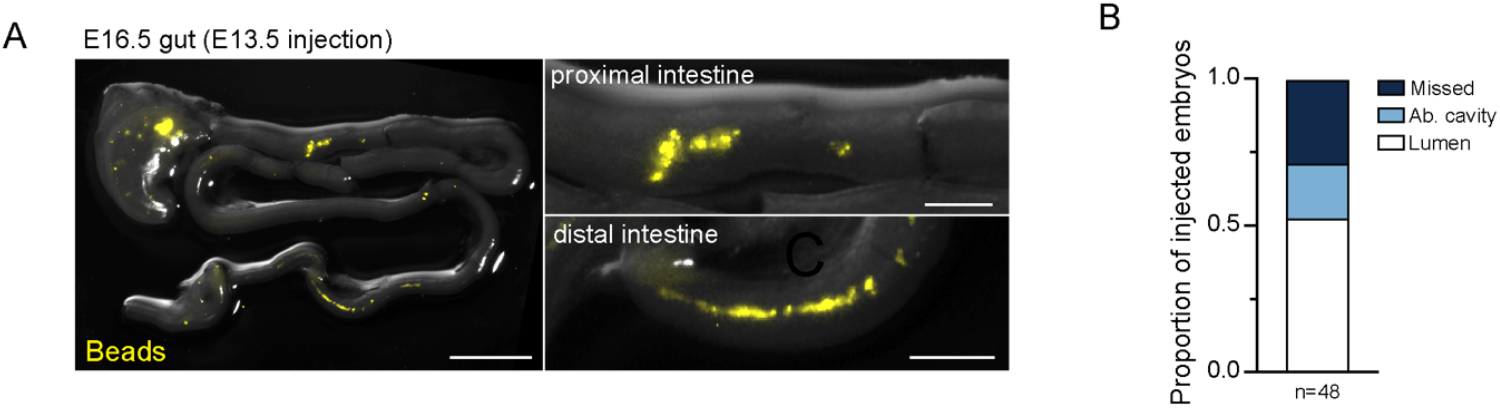
Ultrasound guided, *in utero* microinjection delivery of rabies virus into the embryonic intestinal lumen. (A) E16.5 gut injected at E13.5 with modified rabies virus and fluorescent beads (yellow), confirming luminal injection of the rabies virus. (B) Quantification of luminal injection rates based on the detection of fluorescent beads in tissue sites. Data include 48 embryos from >10 litters, injected at E13.5 or E15.5 and harvested at E16.5 or E18.5. *n* indicates number of embryos. Scale bars, 2mm (A, left panel), 500 µm (A, right panels).

**Fig. S4.**
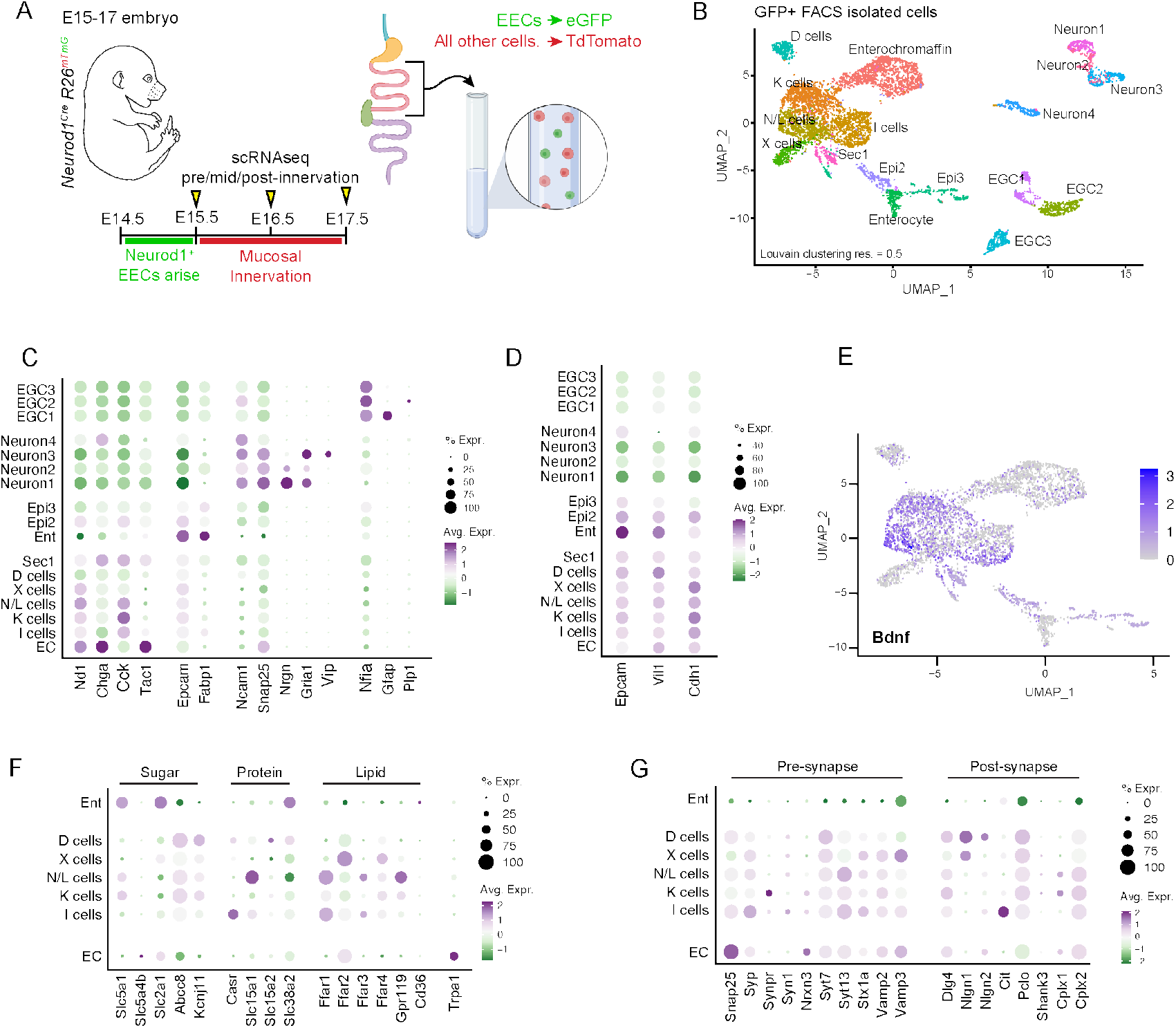
Embryonic EECs are transcriptionally mature. (A) Schematic overview of transgenic labeling approach for FACS-based enrichment of embryonic EECs for scRNA-seq. (B) Uniform manifold approximation and projection (UMAP) plot of isolated cells from E15, E16, & E17 *Nd1*^*mGFP*^ proximal intestine, sorted into 17 distinct clusters. Cells were isolated from 11 individual *Nd1*^*mGFP*^ embryos (three E15.5, four E16.5, and four E17.5). (C) Dot plot of cell type marker genes across all clusters. Here and in all dot plots, green hues represent low expression and purple hues high expression, while the size of the dot indicates the proportion of positive cells in the cluster. (D) Dot plot for epithelial markers: epithelial cell adhesion molecule (*Epcam*), villin (*Vil1*), and E-cadherin (*Cdh1*). (E) UMAP plot of *Bdnf* expression across epithelial clusters. (F) Dot plot expression profile of macronutrient receptors and transporters across embryonic EEC subtypes. (G) Dot plot expression profile of pre- and post-synaptic markers and secretory machinery across embryonic EEC subtypes. EGC = enteric glial cells, Epi = epithelia, Ent = enterocytes, EC = enterochromaffin cells.

**Fig. S5.**
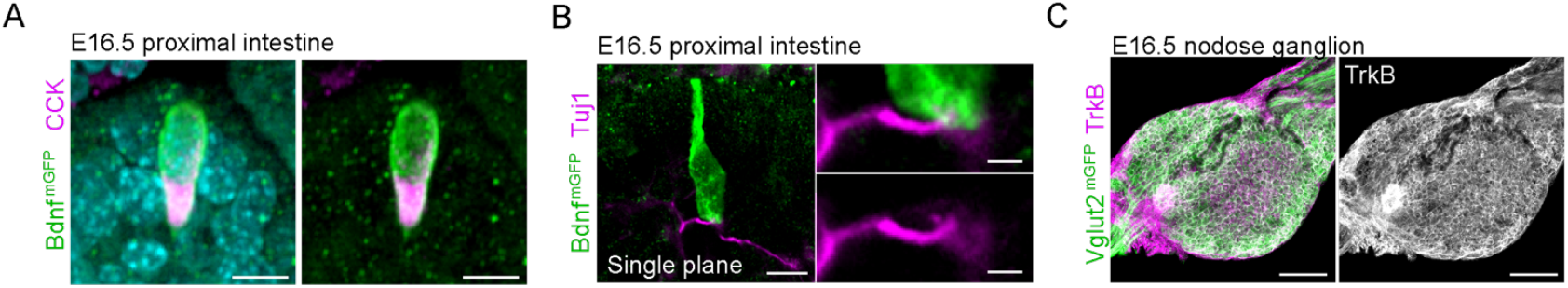
BDNF-TrkB ligand-receptor paired expression in EECs and nodose neurons, respectively. (A,B) Immunofluorescent images of E16.5 proximal intestine from *Bdnf*^*mGFP*^ embryo stained for GFP and CCK (A) or GFP and Tuj1 (B). Right panels in (B) are magnifications of basal contact area showing overlap of EEC mGFP+ membrane and Tuj1+ axon. (C) Max projection image of wholemount nodose ganglion from E16.5 *Vglut2*^*mGFP*^ embryo. Left panel shows mGFP (green) and TrkB (magenta). Right panel shows TrkB (gray) in grayscale. Nearly all nodose neurons are TrkB+. Scale bars, 10 µm (A,B left panel), 2 µm (B right panels), 100 µm (C).

**Fig. S6.**
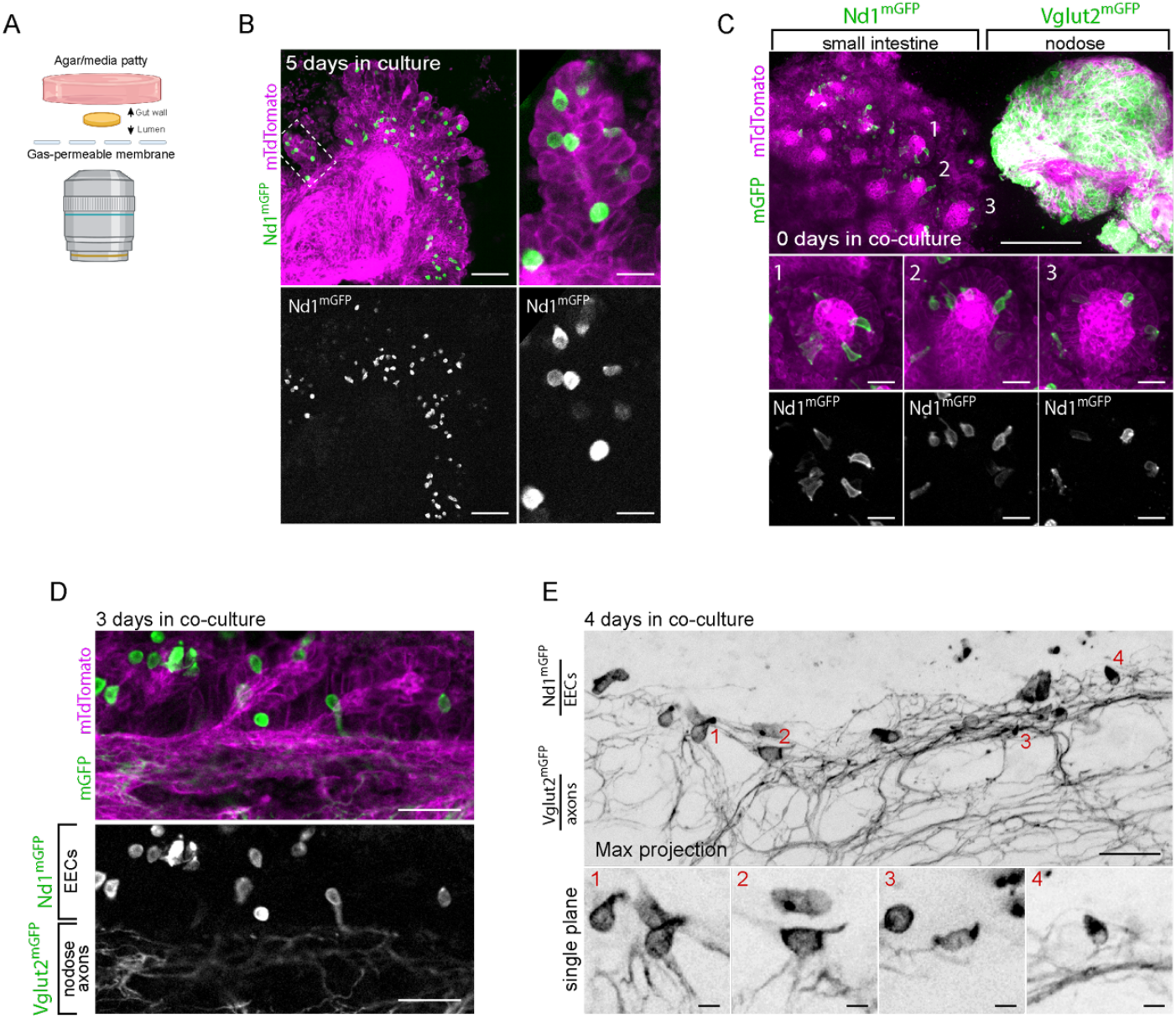
*Ex vivo* co-culture model recapitulates EEC-nodose interactions. (A) Schematic outline of intestinal explant experiment. (B) Still from live imaging of *Nd1*^*mGFP*^ intestine after 5 days of *ex vivo* culture, with mGFP (EECs; green) and mTdTomato (all other cells; magenta). Boxed area of villus shown at higher mag at right. (C) Max projections from freshly plated but-ganglion co-culture (0 day) showing mGFP (green) and mTdTomato (magenta). Numbered insets below are individual villi (numbered in top panel) demonstrating the lack of *Nd1*^*mGFP*^ neuronal fibers in the mucosa. (D) Max projection showing 3-day co-culture where *Vglut2*^*mGFP+*^nodose fibers have approximated villi harboring *Nd1*^*mGFP*^ EECs. (E) Stills from live imaging of 4-day co-culture showing mGFP (black) only. Top image is a max projection. Bottom panels are single plane magnifications of individual EECs in contact with nodose neurons. The red numbers in the bottom panels correspond to the numerically labeled EECs in the top panel. Scale bars, 100 µm (B, left panel), 25 µm (B right panel; C mid and bottom panels), 200 µm (C,E, top panels), 50 µm (D), 10 µm (E, bottom panels).

**Fig. S7.**
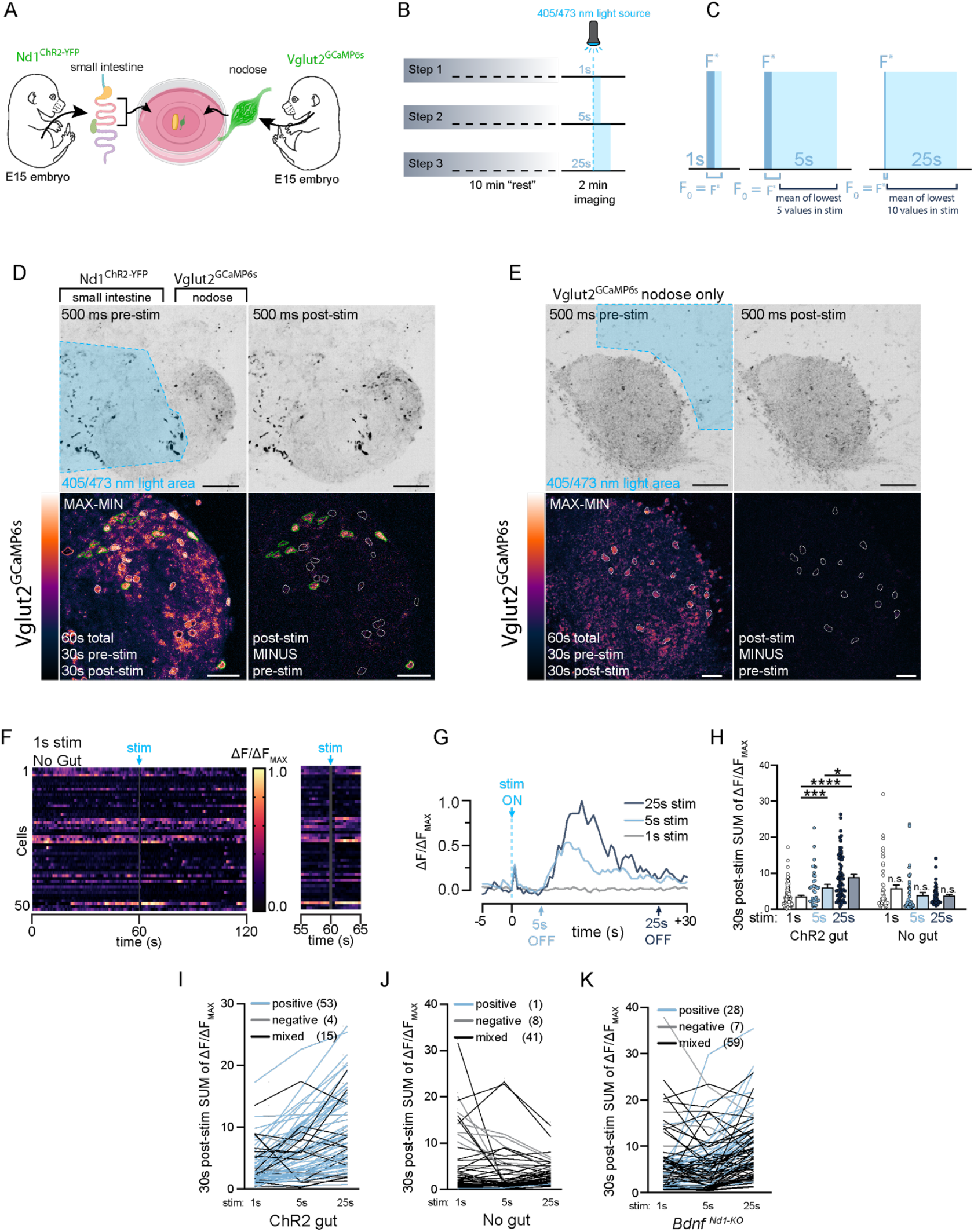
Evoked Ca^2+^ spikes in nodose neurons scale with prolonged EEC optogenetic activation. (A,B) Schematics of gut-nodose explant optogenetic co-culture experiment (A) and intestinal explant optogenetic stimulation protocol (B). (C) Schematic outlining the methodology used to correct for blue-light bleed through during optogenetic stimulation (and quantification of ΔF/Δ F_MAX_). See Materials and Methods for additional details. (D,E) Stills from live imaging of 4 day old optogenetic co-culture of *Nd1*^*ChR2*^ intestine adjacent to a *Vglut2*^*GCaMP6s*^ nodose ganglion (D) and nodose only control (E). Top panels show green fluorescence (YFP in intestinal EECs and GCaMP6s in nodose neurons) immediately prior (left) or immediately after (right) 1s blue light stimulation. Bottom panels show GCaMP6s fluorescence (Gem LUT, see Methods). All cells active during a 1 min period (max-min) are shown at left, while light-activated (post stim minus pre stim) are shown at right. Green outlined cells are optogenetic “responders” while the white outlined cells are “non-responders” who showed spontaneous activity without responding to the optogenetic stimulation. (F) Heatmap kymograph traces of GCaMP6s fluorescence of individual neurons in no gut controls with 1s optogenetic stimulation. Each row represents Ca^2+^ fluctuations for an individual neuron, with 2 minutes of activity shown at left, and 10s of activity— immediately before and after stimulation—shown at right. (G) ΔF-ΔF_MAX_ GCaMP6s fluorescence traces for a single neuron from 4dcc *Nd1*^*ChR2*^ intestine, *Vglut2*^*GCaMP6s*^ nodose co-culture following 1s (gray), 5s (light blue) and 25s (dark blue) stimulation protocols. Note both the immediate spike after the stim as well as the secondary spikes, which scale with the longevity of the stimulation. (H) Quantification of GCaMP6s fluorescence from individual neurons from 4dcc experiments where ΔF-ΔF_MAX_ was summed for the 30s post-stimulation. Each dot represents a single neuron. Note that longer stimulations increase the net GCaMP6s fluorescence in the co-culture experiment but have no effect on the No Gut control. (I-K) Individual cell traces color-coded to reflect the trend in summed (30s SUM of ΔF/ΔF_MAX_) GCaMP6s fluorescence across different stimulation lengths. Light blue lines are cells which displayed a consistent increase in GCaMP6s fluorescence with longer stimulations (positive correlation), while gray lines are cells which showed a consistent decrease (negative correlation). Black lines depict cells with mixed trends. Data shown for *Bdnf*^*WT*^ intestine (I), no gut control (J), and *Bdnf*^*Nd1-cKO*^ co-cultures (K). Numbers of neurons fitting each category are inset in parentheses. Note, most neurons (53/72; 73.6% “positive”) show a positive trend when co-cultured with Nd1ChR2 intestine, while most No gut controls and *Bdnf*^*Nd1-cKO*^ co-cultures show inconsistencies in their response to variable stimulations (No gut: 31/40; 77.5% “mixed”. *Bdnf*^*Nd1-cKO*^: 59/94; 58.5% “mixed”).

**Fig. S8.**
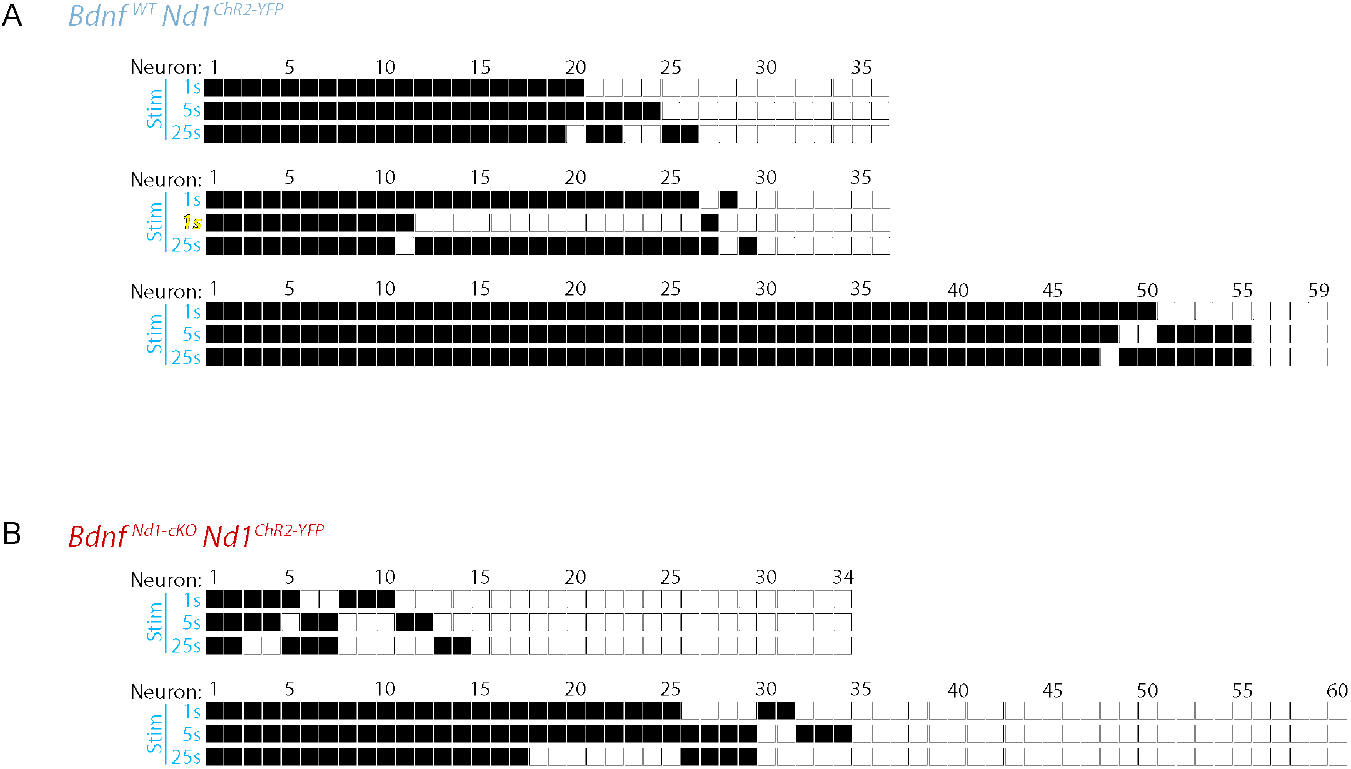
Binary responses of nodose neurons to optogenetic stimulation. (A,B) Immediate response of all nodose neurons from 4 day old co-cultures (4dcc) to optogenetic stimulation. Each column represents a single neuron, while each row reflects the presence (black box) or absence (white box) of an immediate (<1s) evoked spike in GCaMP6s fluorescence (≥0.25 ΔF/ΔF_MAX_) to a 1s (top), 5s (mid), or 25s (bottom) blue-light stimulation. (A) *Bdnf*^*WT*^ *Nd1*^*ChR2-YFP*^ intestinal co-culture results from three separate samples. Note that in the middle sample, the second stimulation was erroneously applied for 1s instead of 5s (yellow text), which may explain the reduction in evoked responses. These data *have not* been excluded from Fig. 4H. (B) *Bdnf*^*Nd1-cKO*^ *Nd1*^*ChR2-YFP*^ intestinal co-culture results from two separate samples.

**Fig. S9.**
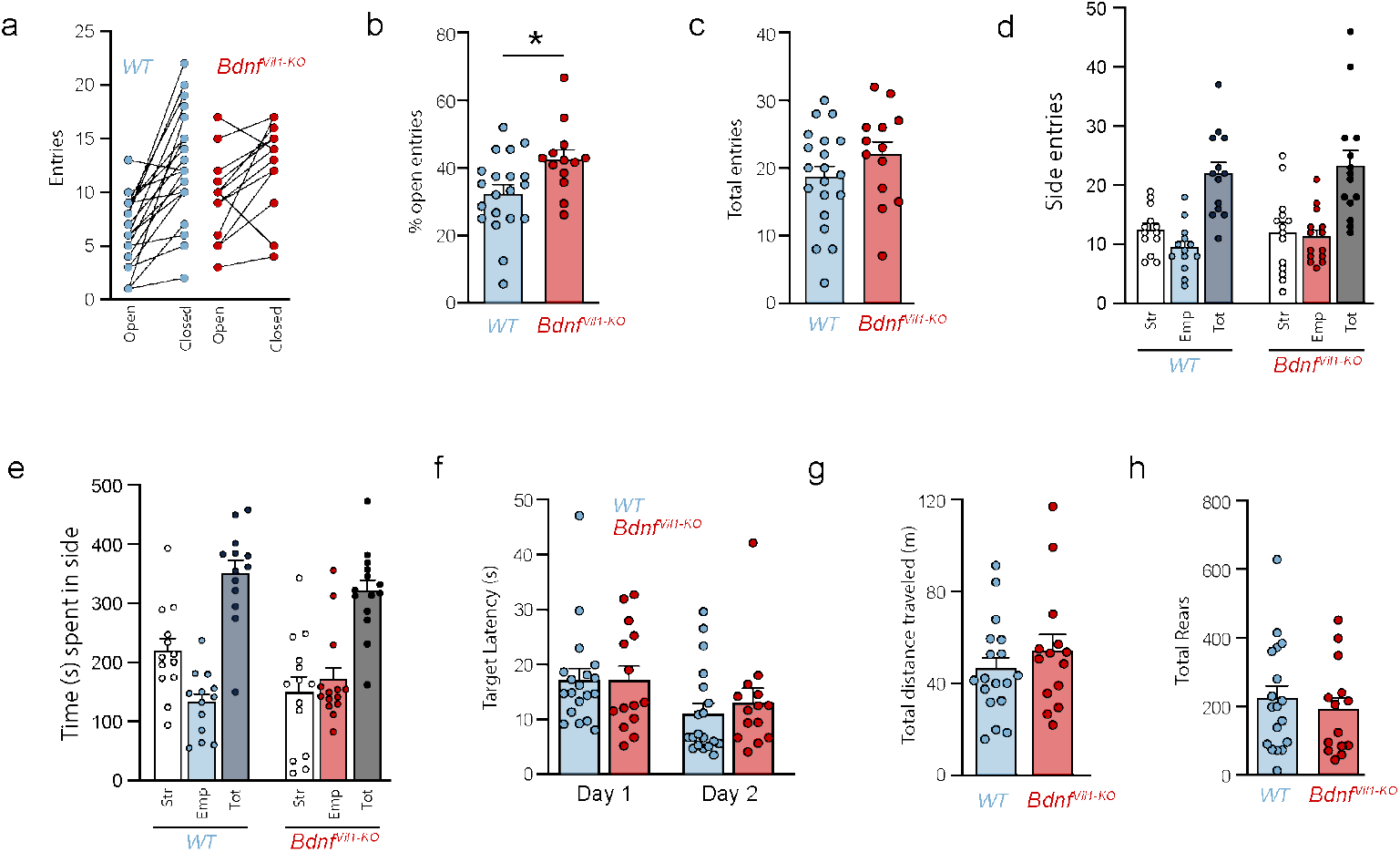
*Villin-Cre* mediated *Bdnf* deletion does not impact general activity or vision. (A) Paired counts of the frequency individual animals entered the open or closed arms of the elevated plus maze. (B) Quantification of the % of open arm entries (over total entries). (C) No significant effects of genotype on total arm entries in the elevated plus maze. (D) Frequency of side entries in the 3-chamber social preference assay, including the stranger chamber (white), chamber with the empty cage (light blue/red), and total side entries (dark blue/black). (E) Summed time (s) spent in the side chambers in the social preference assay. Columns are color-coded as in (D). (F) No significant effects of genotype on latency time (seconds) to reach a visible platform in the Morris Water Maze as a test of visual acuity and swimming ability. (G,H) Open field test, showing no significant effects of genotype on total distance traveled (G) or frequency of rearing movement (H).

**Tabel S1.**
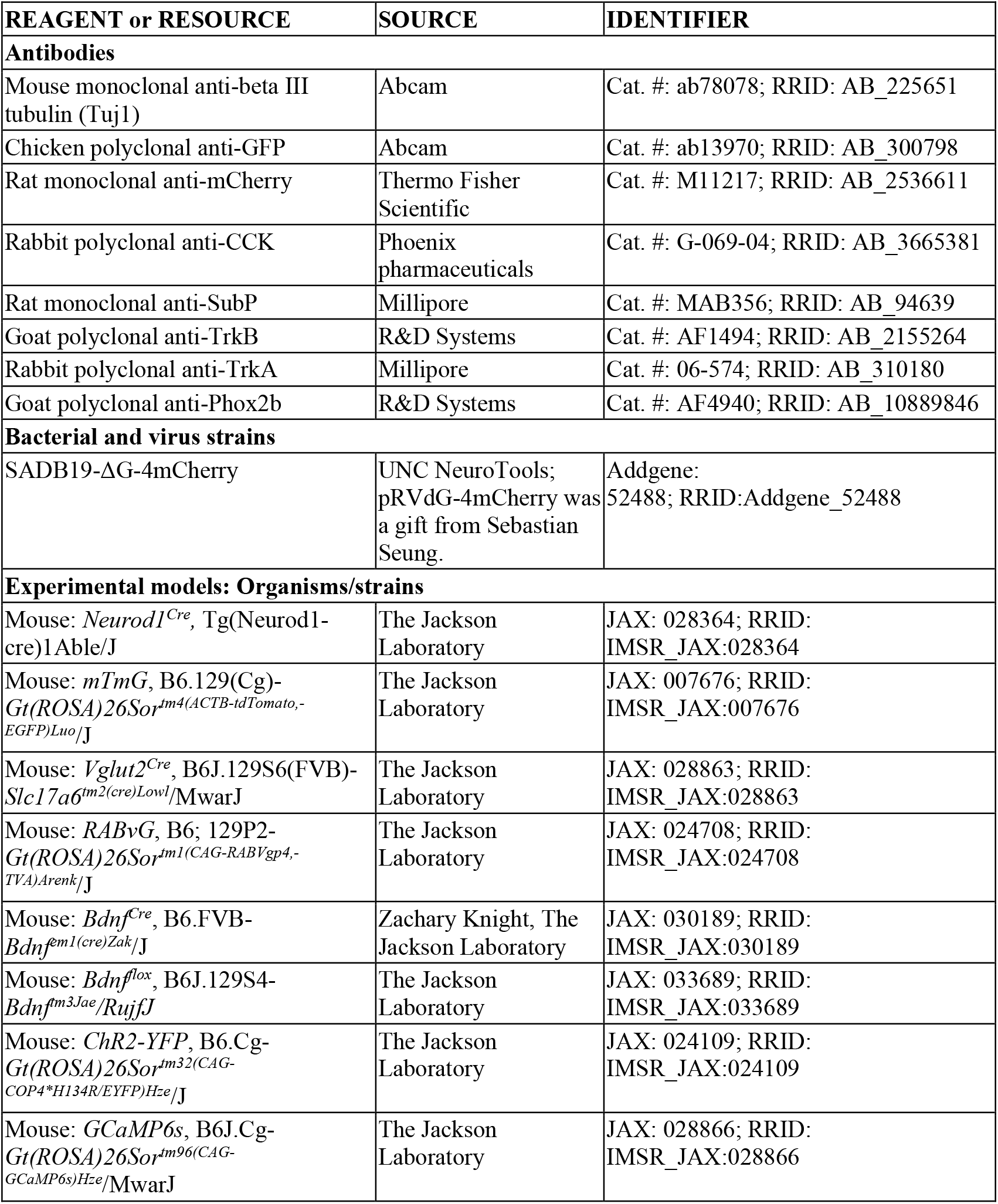
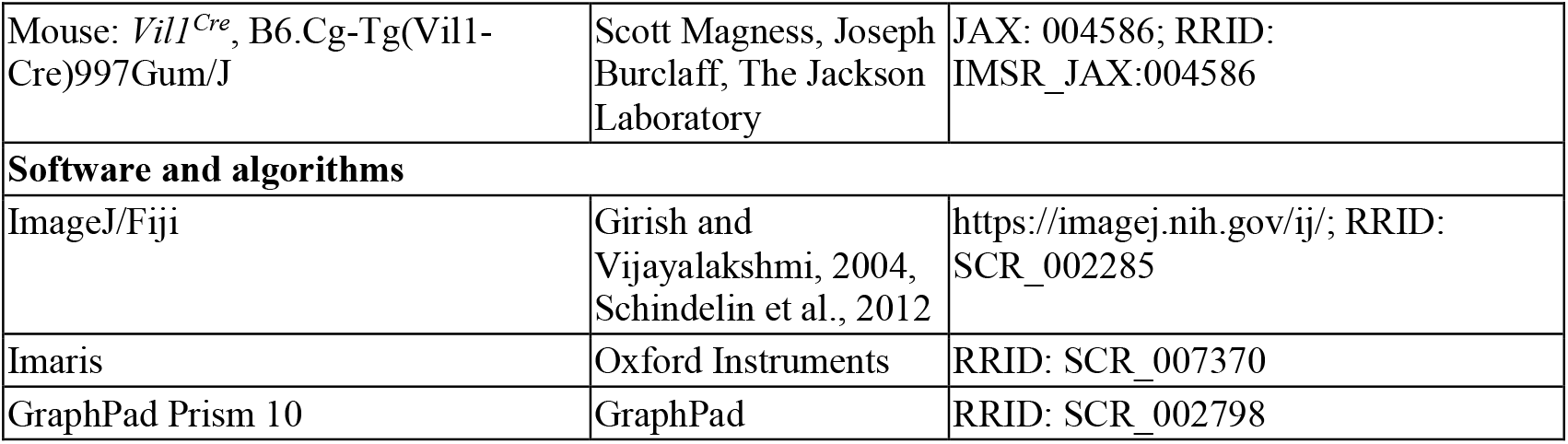
Materials, reagents, and resources table with vendor information. Vendor, catalogue number, and RRID for key resources used in this study. Includes antibodies, viruses, mouse models, and software.

**Movie S1. Newly formed EEC-nodose contact**.

Movie from 1dsc co-cultures of *Nd1*^*mGFP*^ EECs and *Vglut2*^*mGFP*^ projections. Example of an EEC forming a new neuronal contact.

**Movie S2. Lost EEC-nodose contact**.

Movie from 1dsc co-cultures of *Nd1*^*mGFP*^ EECs and *Vglut2*^*mGFP*^ projections. Example of an EEC losing neuronal contact.

**Movie S3. Maintained EEC-nodose contact**.

Movie from 1dsc co-cultures of *Nd1*^*mGFP*^ EECs and *Vglut2*^*mGFP*^ projections. Example of an EEC maintaining neuronal contact.

**Movie S4. “No new” EEC-nodose contacts**.

Movie from 1dsc co-cultures of *Nd1*^*mGFP*^ EECs and *Vglut2*^*mGFP*^ projections. Example of an EEC which does not form any neuronal contact.

